# Capturing Dynamic Neuronal Responses to Dominant and Subordinate Social Hierarchy Members with catFISH

**DOI:** 10.1101/2024.12.19.629477

**Authors:** Madeleine F. Dwortz, James P. Curley

## Abstract

Dominance hierarchies are key to social organization in group-living species, requiring individuals to recognize their own and others’ ranks. This is particularly complex for intermediate-ranking animals, who navigate interactions with higher- and lower-ranking individuals. Using in situ hybridization, we examined how the brains of intermediate-ranked mice in hierarchies respond to dominant and subordinate stimuli by labeling activity-induced immediate early genes and neuronal markers. We show that distinct neuronal populations in the amygdala and hippocampus respond differentially across social contexts. In the basolateral amygdala, glutamatergic Slc17a7+ neurons, particularly dopamine-receptive Slc17a7+Drd1+ neurons, show elevated IEG expression in response to social stimuli, with a higher response to dominant over subordinate animals. Similar patterns are observed among Slc17a7+Oxtr+ neurons in the dorsal endopiriform nucleus and GABAergic Slc32a+ neurons in the medial amygdala. We also identified distinct neural ensembles selectively active in response to dominant and subordinate hierarchy members. We find a higher degree of reactivation among Slc17a7+Oxtr+ ensembles in the dorsal endopiriform nucleus in animals repeatedly presented with the same hierarchy member, as opposed to those presented with a dominant and subordinate member. We observe a similar pattern among Oxtr+ neurons in the dentate gyrus hilus, while the inverse is observed among Slc17a7+ Avrp1b+Oxtr+ neurons in the distal CA2CA3 region. Collectively, our findings reveal how social context is associated with activity changes in social, olfactory, and memory systems in the brain at the neuronal cell type level. This work lays the foundation for further precise cell-type investigation into how the brain processes social information.

## Introduction

A dominance hierarchy is an essential form of social organization that controls resource allocation in social species. The stability of dominance hierarchies and the survival of individuals within them relies on comprehension of hierarchical position, or social status, and the expression of status-appropriate social behavior (Taborsky and Oliveira 2012). This is no simple task, especially for hierarchy-members of intermediate rank, who constantly encounter conspecifics of varying social status and must flexibly adapt their behavior towards animals of relatively higher and lower rank on an ongoing basis. Previous work investigating social status representation in the brain has repeatedly implicated sensory, emotional, and social memory networks that hinge upon the amygdala and hippocampus (Dwortz et al. 2022). Work from our lab demonstrated that the immediate early gene (IEG) *c-fos* shows elevated immunoreactivity in the posteroventral region of the amygdala (MeApv) in response to socially salient cues (Lee et al. 2021). Specifically, this region responds robustly to dominant male urine in both dominant and subordinate mice, regardless of cue familiarity (from an animal of the same social group or novel). Additionally, hippocampal circuits supporting social memory that likely facilitate hierarchy member recognition have been thoroughly characterized (Phillips, Robinson, and Pozzo-Miller 2019; Okuyama et al. 2016; Meira et al. 2018). However, it is still largely unknown which neuronal subpopulations within these regions are differentially responsive to dominant versus subordinate social cues.

To capture how the brains of animals adapt to changes in social context, we employed an innovative IEG-based method, *Arc/H1a* catFISH (cellular analysis of temporal activity by fluorescence in situ hybridization) (Vazdarjanova et al. 2002). This method allowed us to label the responses of IEGs to multiple stimuli and examine how an individual brain reacts to different social contexts in an ethologically relevant manner. IEGs are a class of genes that are rapidly and transiently transcribed upon cell membrane depolarization and labeling of IEGs therefore serves as a proxy of neural activation in response to status-associated stimuli. We selected mid-ranked subject male mice from dominance hierarchies formed in the laboratory and presented them with stimulus pairs consisting of dominant and subordinate members of their hierarchy (**Figure 1A**). Stimulus presentations were confined to a brief period when early investigative sniffing occurs prior to the onset of agonistic behavior. Thus, IEG expression induced by these stimulus events reflects the initial evaluation and perception of social context.

**Figure 1:**
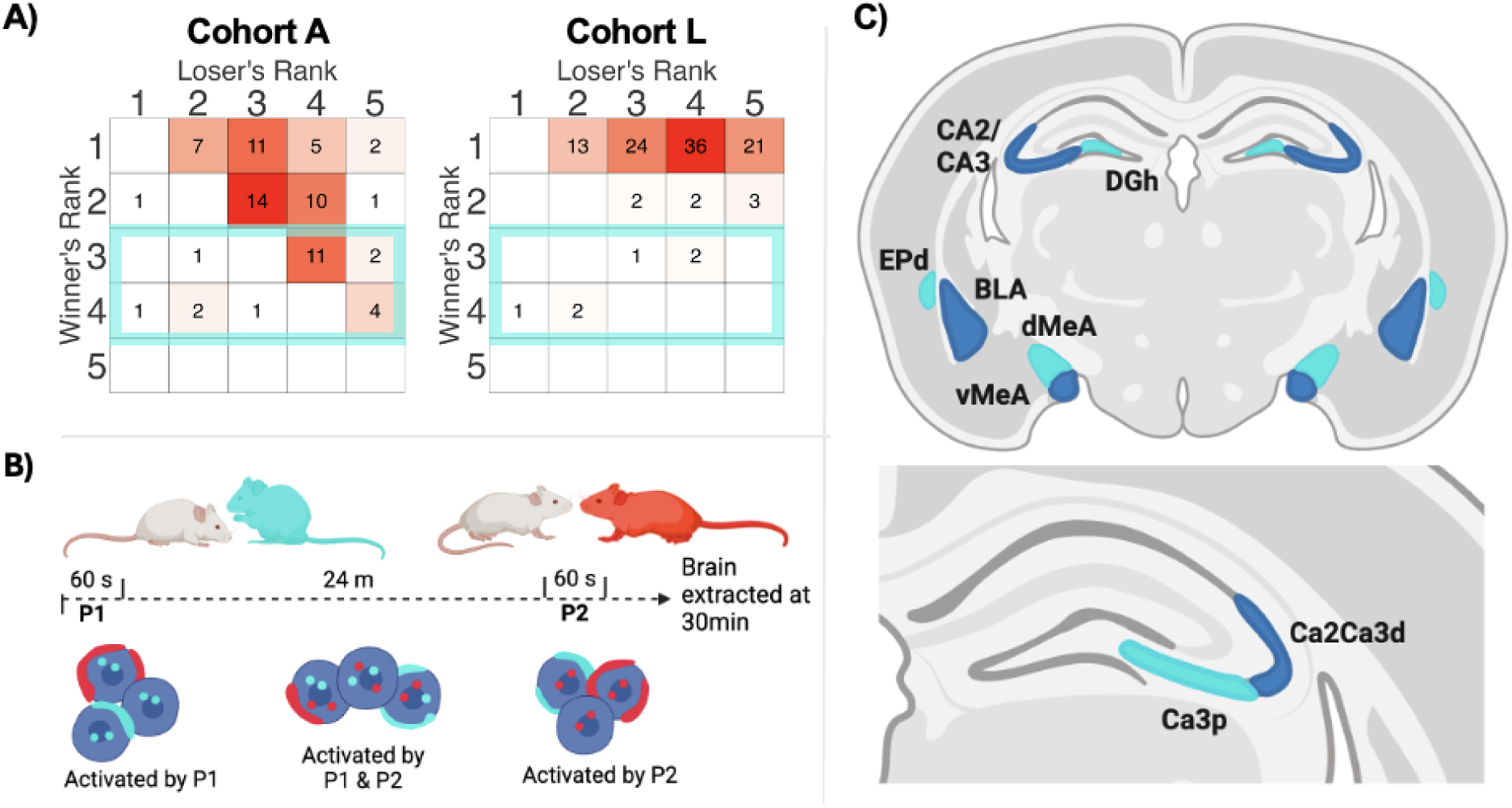
Experimental Overview. **A)** Example sociomatrices show the total frequency of agonistic interactions that occurred between all pairs of individuals in cohort A and cohort L over the entire observation period. Winners of each contest are listed in rows, and losers are listed in columns. Ranks were calculated using the DS method (see methods). Cells of each matrix are colored on a gradient from white (lowest value in each matrix) to red (highest value in each matrix). **B)** Schematic timeline showing order of stimulus presentations and corresponding IEG expression within nuclei. Intranuclear *H1a* and *Arc* are shown in cyan and red, respectively. **C)** Coronal brain slices showing targeted regions (approximately slide 74 of the Allen Mouse Brain Atlas)

Developed originally by Guzowski et al., catFISH labels IEGs with distinct transcription speeds, allowing us to compare neural ensembles activated by two distinct stimulus events separated by approximately 30 minutes (Guzowski et al. 2001). *Arc/H1a* catFISH exploits the different structures of the primary transcripts of IEGs *Arc* and *Homer1a* (*H1a*). While *Arc* mRNA is derived from a short primary transcript, *H1a* mRNA is generated from a longer, more complex primary transcript, requiring additional time for transcription (Kovács 2008). Thus, *Arc* intranuclear foci (INF) appear within approximately 5 minutes of cell activation, while *H1a* INF appear within approximately 30 minutes (**Figure 1B**). Both *Arc* and *H1a* are transcribed upon calcium influx into the postsynaptic membrane, and both play roles in synaptic plasticity by regulating molecular processes involved in synaptic remodeling. Arc is known to modify spine morphology in active dendrites, and H1a has been shown to interact with proteins within the postsynaptic density, including excitatory metabotropic glutamate receptors (Xiao et al. 1998; Vazdarjanova et al. 2002). The role of H1a at inhibitory synapses remains less understood; however, expression of both H1a and Arc have been identified in GABAergic neurons, particularly in regions where these are the principal neurons (Lucas et al. 2008; Vazdarjanova et al. 2006; Imamura et al. 2011). H1a and Arc’s mRNA expression in multiple cell types, their non-overlapping transcription timelines, and transient occurrence within the cell nucleus made these two IEGs suitable targets for the present experiment.

In this study, we examined subregions of the amygdala and hippocampus to assess activation levels and the potential for population-based coding patterns in response to social status cues. We specifically targeted the medial amygdala (MeA), including the MeApv, posterodorsal (MeApd) and anteroventral (MeAav) subregions. We also examined the basolateral amygdala (BLA) and the endopiriform nucleus (EPd). Additionally, we examined the dorsal hippocampus, including pyramidal cell layers of CA3p, distal CA2CA3 (CA2CA3d), and the hilus of the dentate gyrus (DGh) (see **Figure 1C**). We also sought to characterize neuronal populations in these regions based on excitatory and inhibitory properties and their receptivity to neurotransmitters involved in social processing. Identifying excitatory and inhibitory cell types is essential for understanding circuit function, as it reveals whether neurons primarily amplify or suppress activity in their targets. This is especially important for interpreting activity patterns in the amygdala, which contains a heterogeneous array of cell types (Swanson and Petrovich 1998; Yu et al. 2023; Hochgerner et al. 2023). We also targeted receptors heavily involved in modulating social cognition and behavior. Oxytocin receptor expression in the hippocampus and MeA is required for social memory (Raam et al. 2017; Lukas et al. 2013; Ferguson et al. 2001) and, more specifically, the encoding and recognition of social cues (Yao et al. 2017; Li et al. 2017). Vasopressin receptors are also crucial for social memory in the MeA and hippocampus, respectively (Dantzer et al. 1987; Bluthe, Gheusi, and Dantzer 1993; Albers 2012; Wang et al. 2013; Cymerblit-Sabba et al. 2023), and mediate social approach behaviors (Arakawa, Arakawa, and Deak 2010). Additionally, dopamine receptors (DRD1 and DRD2) play multifaceted roles in attention, motivation, learning, and reward processing, which collectively shape social decision-making by balancing risk-avoidance with impulsive behaviors (Homberg et al. 2016; Yamaguchi et al. 2017; Kim et al. 2018; Golden et al. 2019; Tickerhoof et al. 2020). Genes targeted in this study and their protein products include: excitatory and inhibitory markers *Slc17a7*/vesicular glutamate transporter and *Slc32a1*/vesicular GABA transporter, neuropeptide receptors *Oxtr*/oxytocin receptor, *Avrp1a*/vasopressin receptor 1A, and *Avrp1b*/vasopressin receptor 1B, and neuromodulators *Drd1*/dopamine receptor D1 and *Drd2*/dopamine receptor D2. We used RNAscope technology (ACD bio) to manufacture probes and precisely label all nine gene targets.

For each brain region, we identified groups of cells based on their expression of the seven non-IEG mRNAs. We then examined IEG expression levels in each group in response to hierarchy members or an empty cage control. We show that the amygdala and EPd are highly receptive to social stimuli and that specific neural populations exhibit heightened sensitivity to social status. Specifically, *Slc17a7+Drd1+* BLA neurons and *Oxtr+ Slc17a7+* EPd and *Slc32a+* MeA neurons display the highest levels of IEG expression in response to a dominant stimulus. We also assessed overlap, or the reactivation of cells, in response to stimulus pairs. We find that activated *Slc17a7+Oxtr+* ensembles in the EPd exhibit reduced cell re-activation in response to discordant, or non-matching, stimulus pairs (e.g., a dominant animal paired with a subordinate animal), while overlapping ensembles respond to the same dominant or subordinate individual. We identified a similar overlap pattern among DGh *Drd2*+ neurons. Our findings provide methodological insights and enhance our knowledge of how sensory and memory-related systems guide individuals through the complex social environments of dominance hierarchies.

## Methods

### Subjects and housing

Subjects were adult (7-8 weeks) male CD-1 mice (*Mus musculus domesticus*) (Charles River, Houston, TX). Throughout the study, food and water were provided *ad libitum*. Upon arrival, mice were marked for identification using blue non-toxic markers (Stoelting Co.) and pair-housed in standard cages with pine shaving bedding. After three days of habituation to our facility, mice were placed into group housing consisting of two standard-sized rat cages (dimensions: 35.5 x 20.5 x 14cm) separated by two acrylic tubes (diameter: 5cm) with a layer of pine-shaving bedding (**Supplemental Figure S1A**). Subjects remained in group housing for 12-16 days before stimulus presentations and sacrifice. All procedures were conducted with approval from the University of Texas at Austin Institutional Animal Care and Use Committee (IACUC).

### Behavioral observations and dominance analysis

During the first three days of pair housing, experimenters monitored behavior for 1 hour daily and recorded the winners and losers of aggressive interactions. Pairs were then allocated into groups of five so that dominant aggressors were evenly distributed. Groups were observed for no less than one hour every other day for the remainder of the study. During observation periods, observers used all-occurrence sampling to document fighting, chasing, mounting, submissive postures, and fleeing behaviors, noting winners and losers (see **Supplemental Table S1** for ethogram). Behavioral data analysis was then conducted in R Studio using the *compete* package (Curley 2016). The cumulative wins and losses for each individual over the housing period were compiled into frequency win/loss sociomatrices for each cohort (**Figure 1A**, **Supplemental Figure S2**). From these sociomatrices, we calculated the Directional Consistency (DC) of dominance interactions and David’s Scores (DS). In brief, DC assesses the degree to which all agonistic interactions in a group occur in the direction of the more dominant individual to the more subordinate individual within each relationship. It is equal to (H - L)/(H + L) where H is the frequency of behaviors occurring in the most frequent direction and L is the frequency of behaviors occurring in the least frequent direction within each relationship. The significance of DC values were evaluated using a randomization test (Leiva, Solanas, and Salafranca 2008). DS provides an individual dominance rating and ranking for each individual in a group, determining the overall success of each individual at winning contests relative to the success of all other individuals. Briefly, it is derived from the proportion of wins and losses of each individual and is corrected for the frequency of agonistic interactions (de Vries 1995). DS above 0 typically reflects that an animal is socially dominant whereas a DS below 0 typically indicates that an animal is socially subordinate. The most dominant and subordinate individuals in each hierarchy had the highest and lowest DS, respectively. These individuals served as stimulus animals. Subjects were selected from mid-ranked individuals with the third and fourth-highest DS. In total, we used 34 subjects from 23 different hierarchically organized groups.

### Stimulus presentations

Social stimulus presentations occurred in an acrylic chamber (14.9 cm x 16.8cm x 17.75cm) with a thin layer of pine shaving bedding. A removable opaque partition could be placed in the middle of the chamber, dividing it into two compartments (**Supplemental Figure S1B**). For five consecutive days before stimulus presentations, subjects were transferred to a testing room where they acclimated to the entire testing chamber for 15 minutes, followed by an additional 15 minutes of acclimation to each compartment after lowering the partition.

To capture changes in IEG expression in response to different hierarchy members, subjects were exposed to one of five stimulus conditions consisting of paired stimuli in a specific order: 1) dominant and subordinate (Dom-Sub), 2) subordinate and dominant (Sub-Dom), 3) dominant and dominant (Dom-Dom), 4) subordinate and subordinate (Sub-Sub), 5) empty cage and empty cage (EC-EC). The EC-EC condition served as a negative control, providing a baseline measurement of brain activity induced by experimental handling. Stimulus presentations proceeded as follows. Subjects first habituated to the whole chamber for 30 minutes, followed by the front compartment for 30 minutes. Then, the first stimulus presentation (P1) occurred. Subjects were presented with a stimulus animal or a control stimulus where the experimenter opened the chamber and briefly placed their hand inside. A social stimulus consisted of placing the most dominant or subordinate individual from the subject’s home cage into the compartment with the subject. Both mice were allowed to investigate each other for no more than 60 seconds, a brief duration that mitigated fighting. The experimenter closely monitored social interactions and removed the stimulus mouse before 60 seconds if individuals were postured to fight. Warning signs of impending fights included progressively increased sniffing involving physical contact and tail rattling. Trials that involved fighting were removed from the experiment, resulting in the following number of subjects in each stimulus presentation condition (Dom-Sub: n=7, Sub-Dom: n=6, Dom-Dom: n=8, Sub-Sub: n=7, EC-EC: n=9). After removing the stimulus mouse, the partition was briefly lifted, allowing the subject to move into the back compartment. This design mitigated the carry-over of bedding and social odors the stimulus mouse could leave behind. Then, 25 minutes after the onset of P1, the second stimulus presentation (P2) occurred. For social stimulus presentations, a second stimulus mouse from the subject’s home cage was placed in the compartment with the subject mouse. This individual was either the same mouse from the first presentation or a hierarchy member of the opposing social status. Then, 5 minutes after the onset of P2, the subject was sacrificed. This stimulus presentation timeline facilitated the capture of nuclear *H1a* and *Arc* expression induced by P1 and P2, respectively (**Figure 1B**).

All stimulus presentations were video recorded from above, and sniffing behavior was manually coded using BORIS software (Friard and Gamba 2016). Observers recorded stimulus presentation onset when stimulus animals entered the chamber and offset when they were removed. They also recorded “sniffing” when the subject’s nose was in contact with or came close to the stimulus animal. The location of sniffing (e.g., perioral or anogenital regions) was also recorded to account for potential differences in olfactory inputs across subjects. Statistical analysis of stimulus presentation duration and time spent sniffing the stimulus animal, or sniffing duration, across stimulus conditions was conducted in R Studio with the package *lme4* (R Core Team 2003; Bates et al. 2015). (R Core Team 2003; Bates et al. 2015). We fit general linear mixed effect models (GLMMs) to assess whether presentation and sniffing duration changed as a function of the stimulus presentation order (P1 or P2) or the stimulus animal’s status (dominant or subordinate).

### Tissue preparation

Immediately upon sacrifice, brain tissue was flash-frozen in dry ice-cooled hexanes and stored at −80°F until further processing. Brain tissue was then transferred to a −20°C cryostat chamber (Leica Microsystems), where it incubated for approximately 30 minutes before sectioning. We used the publicly available Allen Brain Atlas for mice to identify and section regions of interest (approximately slide 74 of the atlas, see **Figure 1C**). Two 20μm thick sections were then made onto SuperFrost Plus charged slides. Sections were briefly thawed to adhere to the slide but were immediately returned to the −20°C cryostat chamber until sectioning was completed. Slides were then placed in a slide box and inside an air-tight plastic zip-lock bag and stored at −80°F until further processing.

### RNAscope in situ hybridization and imaging

RNAscope in situ hybridization (*ISH*) Hiplex v2 assay was performed according to the Advanced Cell Diagnostics (ACD Bio) protocol for fresh frozen tissue. We used positive and negative control probes supplied by the kit (ACD Cat. No. 320881 and 320871, respectively) to verify RNA integrity and binding specificity. In addition to a DAPI stain for nuclei, three fluorophores (FITC, Texas Red, and Cy5) were used in three iterative binding and cleaving rounds to label nine target genes (**Supplemental Figure S3**). The following probes from ACD Bio were used: Mm-Homer1-O2-T1 (*H1a*), Mm-Arc-T2 (*Arc*), Mm-Oxtr-T3 (*Oxtr*), Mm-Slc17a7-T5 (*Slc17a7*), Mm-Avpr1a-T6 (*Avrp1a*), Mm-Avpr1b-T7 (*Avrp1b*), Mm-Slc32a1-T9 (*Slc32a1*), Mm-Drd1a-T10 (*Drd1*), and Mm-Drd2-T11 (*Drd2*). After each round, image scans were taken of the amygdala and hippocampus in the right and left hemispheres at 20X magnification using a Leica DMi8 inverted microscope and LASX software (Leica Microsystems). At least two brain sections were imaged for each subject. Only high-quality tissue sections were counted. In cases where a brain region’s anatomy was ambiguous, or there was tissue damage, additional sections were processed. As a result, the number of image scans per brain region per subject ranged from one to four (mean = 3 images). Images from each round were overlayed using the DAPI signal as a reference with ACD RNAscope HiPlex Image Registration Software. The resulting 10-channel image (one DAPI channel and a channel for each of the nine probes) was imported into the LASX software for semi-automated quantification.

### Quantification of ISH signals

The LASX 2D Analysis Software (Leica Microsystems) was used to remove background signals, count cells, and quantify the presence of the nine probes in each cell. The following brain regions were manually outlined using the Allen Brain Atlas for mice as a reference: BLA, MeApd, MeApv, MeAav, EPd, CA2CA3d, CA3p, and DGh. For channels corresponding to *H1a* and *Arc*, conservative thresholding was used to eliminate diffuse cytoplasmic signals and preserve intense intra-nuclear foci (INF). Background signal was also eliminated for all channels to reduce false positives, resulting in some unlabeled cells. A separate report was then generated for each ROI. Each report consists of a results table where each row represents a DAPI-stained nucleus, and each column represents one of the nine probes and whether a signal was detected “inside” that nucleus or “touching” the nucleus’s border in the cell’s cytoplasm.

### Statistical analysis of ISH data

We analyzed stimulus-associated IEG expression in distinct cell populations for each brain region. First, we classified the presence of mRNA targets in each cell. As we aimed to count *H1a* and *Arc* INF, these IEGs were considered present in a given cell if signals were detected inside the DAPI-stained nucleus. Each of the seven non-IEG targets was considered present if a signal existed in the nucleus or cytoplasm. Counts across both hemispheres and sections for a given mouse were aggregated and normalized by the total number of cells.

We identified unique cell populations based on the labeling of the non-IEG targets. To remove potential artifacts, we applied a filtering step to remove gene combinations for which less than 3% of cells were labeled in a given brain region. Statistical analyses were then applied to the proportions of IEG-positive cells in these cell populations. We used the *lme4* package and fit GLMMs for binomially distributed data to proportions of IEG-expressing cells. We included the number of cells in each cell type as weights in the model. This accounted for variation in the number of cells in each subject and ensured that we did not conflate more activation of a given cell population with a greater abundance of that population. We also included Subject ID and batch as random factors in the models. We applied two primary models, one in which social context (i.e. presence or absence of social stimulus) was a fixed factor, and a second in which the type of stimulus (i.e., dominant, subordinate, or empty cage) was a fixed factor.

We also employed more complex models to examine associations between IEG expression and non-central aspects of stimulus presentations, such as the presentation order and the specific IEG (P1/*H1a* or P2/*Arc*), as well as presentation duration and time spent sniffing stimulus animals. Subjects in the EC-EC condition were not included in the presentation and sniffing duration analyses. An optimal model was chosen for each brain region and cell population based on *Akaike* information criterion (*AIC*), a measure that aids in the selection of the most parsimonious model that best fits the data (see **Supplemental Table S2** for list of cell populations and the best-fit models). Pairwise posthoc testing with False Discovery Rate (FDR) correction for multiple hypothesis testing was performed on all model results to evaluate parameter comparisons and adjust p values.

We also examined IEG expression overlap, a measurement of how many cells activated in P1 were re-activated by P2. Overlap was calculated for each cell type using the formula below (Saidov 2019).

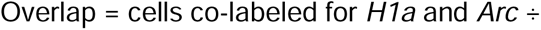

(cells labeled for just *H1a* + cells labeled for just *Arc* - cells co-labeled for *H1a* and *Arc*) We applied GLMMs to test the association between overlap and each stimulus condition, including subject ID and batch as random factors. We compared overlap between each of the five stimulus pairings, as well as between social and non-social stimulus pairings, and concordant (i.e. two identical stimuli) versus discordant stimuli. We analyzed the effects of stimulus concordance across all subjects, including those in the EC-EC condition, as well as those presented with social stimuli. This approach enabled us to compare the impacts of matching versus non-matching stimuli overall, and specifically within the context of social stimuli.

## Results

### Male mice formed significantly linear dominance hierarchies

All 23 social groups of male mice formed linear dominance hierarchies, with a mean DC of 0.933 ± 0.089 (range 0.667 – 1.00) and a mean h’ of 0.806 ± 0.109 range (0.6 – 1.00) (see **Supplemental Table S3** for full dominance results and **Supplemental Figure S2** for sociomatrices).

### The status of stimulus animals did not affect the duration of presentations

We presented mid-ranked individuals with stimulus pairs consisting of the most dominant (Dom) or subordinate (Sub) members from their hierarchy or an empty-cage control (EC). There was not a significant effect of presentation order (P1 or P2) nor the status of the stimulus animal (Dom or Sub) on the duration of the presentation or amount of time subjects spent sniffing stimulus animals (**Supplemental Figure S4, Supplemental Table S4, Supplemental Table S5**). There was a slight trend, although not statistically significant, where sniffing duration in P1 was slightly longer than P2 (P2 – P1: β = −5.388 ± 2.845, p = 0.058). When we assessed within-subject variations in presentation and sniffing durations across stimulus conditions, we observed that subjects presented with Dom-Dom, exhibited less sniffing in P2 (P2 – P1: β = −6.628 ± 1.392, p <0.001), despite the total duration of the presentation period in P2 being slightly longer in these subjects (P2 – P1: β = −5.163 ± 2.10, p = 0.014) (**Supplemental Figure S4, Supplemental Table S5**). In other words, animals exhibited reduced sniffing in their second encounter with a dominant animal despite having more opportunities to sniff.

### IEG expression is associated with social context

We employed RNAscope in situ hybridization to label the IEGs *Arc* and *H1a*, identifying active neuronal populations in the amygdala and hippocampus. Detection of *H1a* and *Arc* INF indicated activation in response to P1 and P2, respectively. We also labeled inhibitory and excitatory markers *Slc32a* and *Slc17a7*, and neurotransmitter receptor genes *Oxtr, Avrp1b, Avrp1a, Drd1*, and *Drd2* to characterize these active populations. Binomial GLMMs were applied to test the association between the stimuli presented and the proportion of IEG-labeled cells to assess putative responses to stimuli.

Our findings show that most principal neuronal populations in the amygdala are more responsive to social stimuli than empty cage controls, with further refined responses to dominant versus subordinate cues. Among all *Slc17a7*+ BLA neurons, we observed elevated responses to social stimuli, with responses being higher to Dom compared to both Sub and EC (Dom > Sub > EC) (Social – EC: β =0.501±0.215, p = 0.033; EC – Dom: β = −0.519 ± 0.215, p = 0.027; Sub – Dom: β = −0.038 ± 0.021, p = 0.111; Sub – EC: β = 0.481 ± 0.215, p = 0.041) (**Figure 2A**). Approximately 16.6 ± 12.3% of *Slc17a7*+ BLA neurons were co-labeled for *Drd1* and 14.3 ± 4.8 % were co-labeled *Oxtr.* The subpopulation of *Slc17a7*+*Oxtr*+ BLA neurons also exhibited elevated responses to social versus non-social stimuli but did not respond significantly more to Dom over Sub (Social – EC: β = 0.544 ± 0.244, p = 0.042; EC – Dom: β = −0.521 ± 0.247, p = 0.056; Sub – Dom: β = 0.051 ± 0.052, p = 0.372; Sub – EC: β = 0.571 ± 0.247, p = 0.035; **Figure 2B**). In contrast, *Slc17a7*+*Drd1+* BLA neurons did not exhibit significantly higher responses to Sub stimuli over EC, but they did show the most pronounced response to Dom over Sub (Social – EC: β = 0.415 ± 0.240, p = 0.118; EC – Dom: β = −0.535 ± 0.240, p = 0.042; Sub – Dom: β = −0.240 ± 0.05, p < 0.001; Sub – EC: β = 0.296 ± 0.240, p = 0.267; **Figure 2C**). Among subjects presented with a combination of Dom and Sub stimuli, this *Slc17a7*+*Drd1+* population exhibited a reduced response to Sub (**Supplemental Figure S5A**).

**Figure 2:**
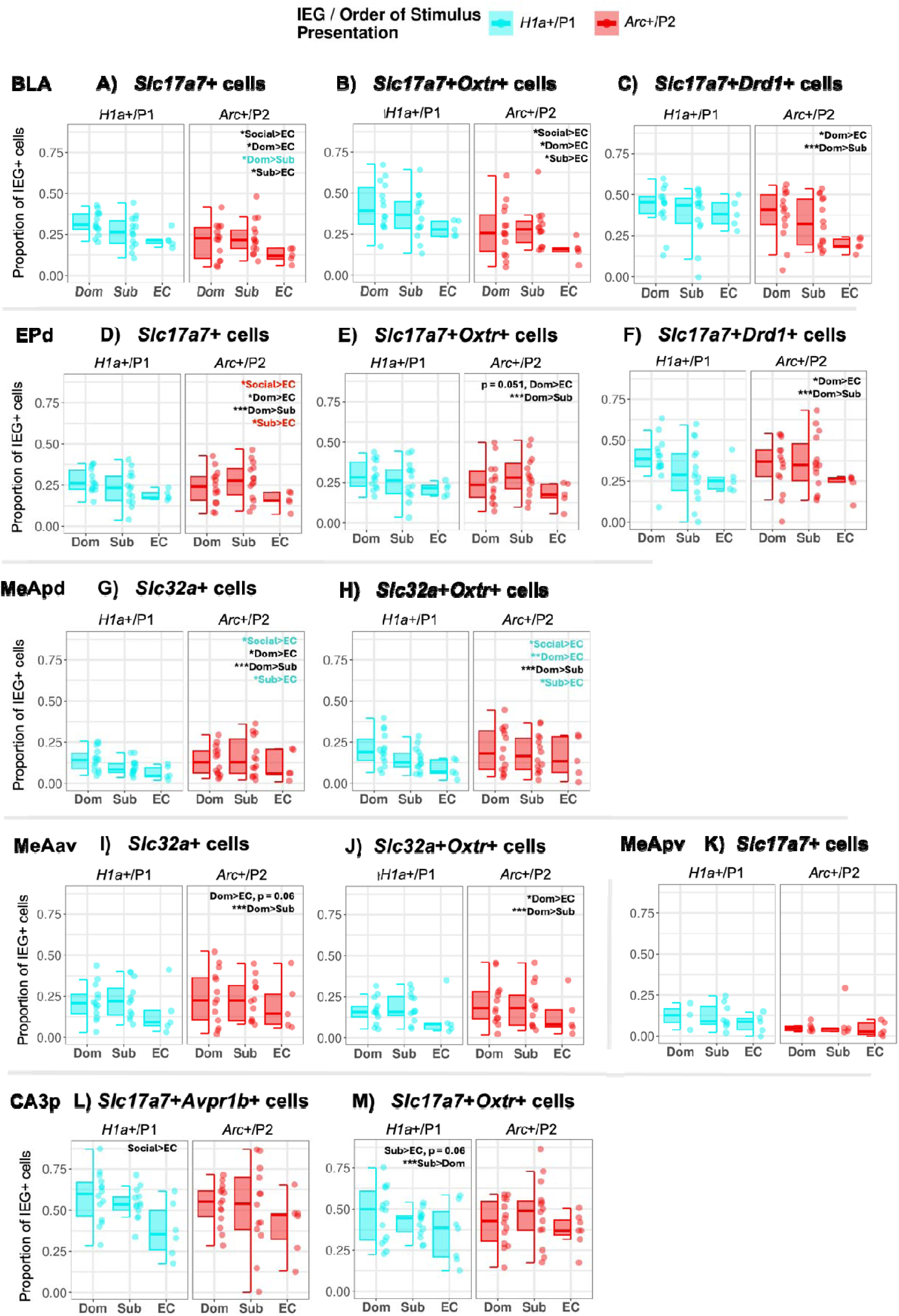
Differential IEG Responses Across Stimuli. Boxplots showing IEG expression in response to Dominant (Dom) and Subordinate (Sub) animals and an empty cage control (EC) for major cell types within each brain region. Responses for each IEG/Presentation order channel are shown separately for visual clarity. Significant comparisons are stated in the upper right of one of the plots (p < 0.05 (*), p < 0.01 (**), p < 0.001 (***)). Black text indicates that the comparison was significant with IEG/Presentation Order included as a random effect in the statistical model. We also examined the interaction of IEG/Presentation Order and stimulus types to determine if differential responses only arose in a specific IEG/Presentation Order channel. Comparisons that were only significant within the h1a/P1 channel are indicated by cyan text. Comparisons that were only significant within the arc/P2 channel are indicated by red text.

We observed a similar response trend in the EPd, where all *Slc17a7*+ neurons collectively displayed stronger responses to Dom compared to both Sub and EC, but responses to Sub were not substantially greater than those to EC. Interestingly, as in the BLA, differential responses between Dom and Sub were most pronounced among *Slc17a7*+*Drd1*+ neurons (*Slc17a7*+: Social – EC: β = 0.448 ± 0.239, p = 0.116; EC – Dom: β = −0.559 ± 0.246, p = 0.058; Sub – Dom: β = −0.226 ± 0.034, p < 0.001; Sub – EC: β = 0.333 ± 0.246, p = 0.243, **Figure 2D**; *Slc17a7*+*Oxtr*+: Social – EC: β = 0.427 ± 0.2588, p = 0.099; EC – Dom: β = −0.734 ± 0.298, p = 0.051; Sub – Dom: β = −0.184 ± 0.046, p < 0.001; Sub – EC: 0.334 ± 0.265, p = 0.207, **Figure 2E**; *Slc17a7*+*Drd1*+: Social – EC: β = 0.492 ± 0.283, p = 0.144; EC – Dom: β = −0.734 ± 0.298, p = 0.037; Sub – Dom: β = −0.496 ± 0.059, p < 0.001; Sub – EC: 0.238 ± 0.298, p = 0.492; **Figure 2F, Supplemental Figure S5B**). The majority of EPd neurons were glutamatergic, with approximately 49.9 ± 7.5% of cells labeled for *Slc17a7*, compared to 6.3 ± 2.1% labeled for GABAergic marker, *Slc32a*. Among *Slc17a7*+ EPd neurons, a large percentage, 51.8 ± 8.4%, were labeled for *Oxtr*, 14.3 ± 7.3% were labeled for *Drd1*, and 15.9 ± 9.2% were labeled for both *Oxtr* and *Drd1*.

Specific GABAergic populations also exhibited heightened responses to Dom stimuli. Principal *Slc32a*+ neurons of the MeApd and MeAav responded more to Dom than Sub and EC (*Slc32a*+ neurons: MeApd: Social – EC: β = 0.609 ± 0.330, p = 0.085; EC – Dom: β =-0.729 ± 0.332, p = 0.041; Sub – Dom: β =-0.252 ± 0.027, p < 0.001; Sub – EC: β = 0.477 ± 0.332, p = 0.178, **Figure 2G**; MeAav: Social – EC: β = 0.622 ± 0.347, p = 0.112; EC – Dom: β −0.750 ± 0.357, p = 0.060; Sub – Dom: β −0.274 ± 0.035, p < 0.001; Sub – EC: β = 0.476 ± 0.358, p = 0.243, **Figure 2I**). This pattern was also observed among *Slc32a*+*Oxtr*+ neurons, specifically (*Slc32a*+Oxtr+ neurons: MeApd: Social – EC: β = 0.586 ± 0.342, p = 0.111; EC – Dom: β = - 0.682 ± 0.340, p = 0.062; Sub – Dom: β = −0.199 ± 0.045, p < 0.001; Sub – EC: β = 0.483 ± 0.340, p = 0.182, **Figure 2H, Supplemental Figure S5C**; MeAav: Social – EC: β = 0.450 ± 0.361, p = 0.275; EC – Dom: β = −0.554 ± 0.372, p = 0.187; Sub – Dom: β = −0.224 ± 0.063, p = 0.001; Sub – EC: β = 0.330 ± 0.372, p = 0.432, **Figure 2J**) Approximately 26.6 ± 7.9% of *Slc32a*+ MeApd neurons were co-labeled for *Oxtr*. More posterior tissue sections captured the MeApv, which, in stark contrast to the MeAav, contains an abundance of *Slc17a7*+ neurons. The heightened response of *Slc32a*+ neurons to dominant stimuli persisted into the MeApv, but notably, no significant response patterns were observed among *Slc17a7*+ neurons in this region (*Slc32a*+: Social – EC: β = 1.126 ± 0.530, p = 0.080; EC – Dom: β = −1.167 ± 0.539, p = 0.073; Sub – Dom: β = −0.076 ± 0.159, p = 0.728; Sub – EC: β = 1.091 ± 0.537, p = 0.095; *Slc17a7*+: Social – EC: β = 0.324 ± 0.412, p = 0.551; EC – Dom: β = −0.285 ± 0.415, p = 0.604; Sub – Dom: β = −0.061 ± 0.089, p = 0.604; Sub – EC: β = 0.346 ± 0.413, p = 0.525, **Figure 2K**). It should be noted that the MeApv dataset was parsed from the MeAav dataset, resulting in fewer samples per stimulus treatment.

In contrast to the amygdala, the hippocampus exhibited a lack of differential activity to Dom and Sub stimuli. One cell population, *Slc17a7*+*Avrp1b*+ neurons in CA3p, responded generally to the presence of a social stimulus without further differentiation between Dom and Sub (Social – EC: β = 0.654 ± 0.250, p = 0.024; EC – Dom: β = −0.606 ± 0.256, p = 0.039; Sub – Dom: β = 0.101 ± 0.063, p = 0.163; Sub – EC: β = 0.707 ± 0.256, p = 0.017, **Figure 2L**). Approximately 10.1 ± 8.5% of all *Slc17a7*+ CA3p neurons were co-labeled for *Avrp1b*+. We also observed a slight increase in response to Sub in *Slc17a7*+*Oxtr*+ CA3p neurons (Social – EC: β = 0.367 ± 0.239, p = 0.180; EC – Dom: β = −0.219 ± 0.247, p = 0.415; Sub – Dom: β = 0.312 ± 0.055, p < 0.001; Sub – EC: β = 0.531 ± 0.247, p = 0.061, **Figure 2M, Supplemental Figure S5D**). Approximately 15.7 ± 6.4% of all *Slc17a7*+ CA3p neurons were co-labeled for *Oxtr*+.

### Stimulus duration shapes IEG expression throughout the brain

We examined the effects of presentation duration among subjects presented with social stimuli. We observed a significant positive association between IEG expression and stimulus presentation duration among most principal neuron types (**Figure 3**). This relationship was most pronounced in the glutamatergic *Slc17a7*+ neurons of the BLA and MeApv (BLA: β = 0.004 ± 0.001, p = 0.004; MeApv: β = 0.027 ± 0.006, p < 0.001) (**Figure 3A,B**). A slight positive association was also noted among *Slc17a7*+ hippocampal neurons (CA3p: β = 0.006 ± 0.002, p = 0.001; CA2CA3d: β = 0.005 ± 0.002, p = 0.033) (**Figure 3E,F**). We also observed this response pattern among *Slc32a*+ MeA neurons; however, there was a significant interaction between presentation duration and stimulus order, such that presentation duration was correlated with *Arc* but not *H1a* expression (P1/*H1a* ∼ presentation duration slope – P2/*Arc* ∼ presentation duration slope: MeApd: β = −0.265 ± 0.018, p < 0.001; MeAav: β = −0.206 ± 0.025, p < 0.001) (**Figure 3C,D**). Note that there also appears to be an interaction between presentation order and duration and among *Slc17a7*+ neurons in the hippocampus (**Figure 3E,F**), such that *H1a* expression appears to be more positively associated with presentation duration. This was not significant due to the inclusion of subject ID as a random effect in the statistical model. In fact, *Arc* expression was slightly more positively associated with presentation duration when we account for subject ID and examine within-subject responses.

**Figure 3:**
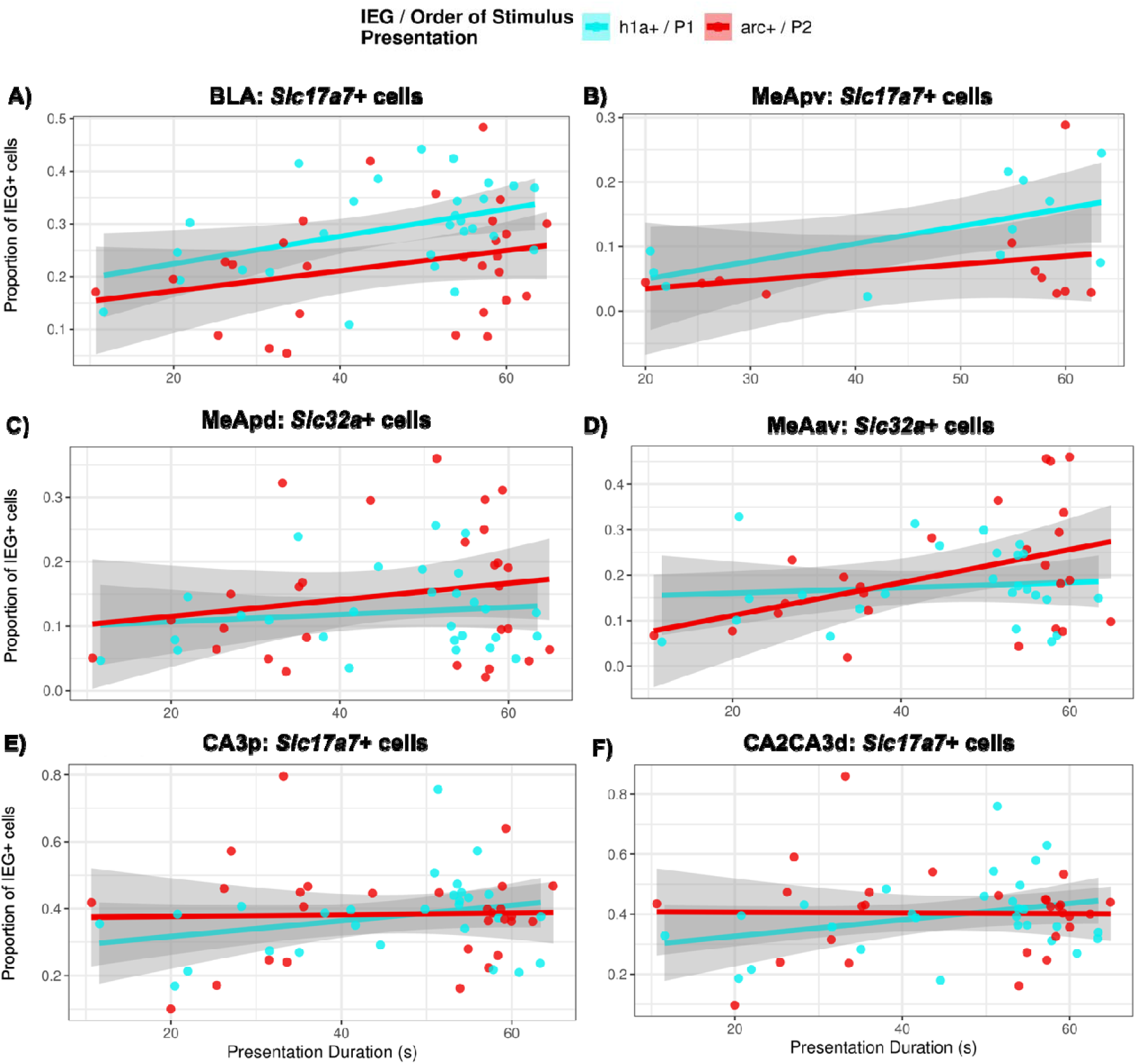
Association Between IEG Expression and Presentation Duration. Significant positive correlations between IEG expression and presentation duration were identified in the BLA (**A**), MeApv (**B**), MeApd (**C**), MeAav (**D**), CA3p (**E**), and CA2CA3d (**F**).

Sniffing duration was mediated by presentation duration; consequently, responses linked to one factor tended to be associated with the other. Incorporating sniffing duration as a fixed factor in the statistical model resulted in a better fit than presentation duration for a limited number of cell populations. Among these, *Slc32a*+*Drd1*+ neurons in the MeAav, which constituted 14.9 ± 8.5% of all Slc32a+ MeAav neurons, demonstrated responses positively associated with sniffing duration (β = 0.015 ± 0.006, p = 0.012).

### Stimulus presentation order effects shape responses to social stimuli

Response patterns observed in the amygdala (Dom > Sub > EC) were more pronounced and, in many cases, only reached statistical significance in a specific order/IEG channel (**Table S6**). The nature of this interaction was not associated with *Slc17a7+* and *Slc32a+* cell types. For example, the differential responses of *Slc17a7+* BLA neurons were more robust with respect to P1/*H1a* (**Figure 2A**). However, the same differential response pattern among *Slc17a7+Drd1+* BLA neurons was more robust with respect to P2/*Arc (***Figure 2C**). In the EPd, *Slc17a7*+ neurons also exhibited significantly higher *Arc*, not *H1a*, expression in response to Dom over EC (**Figure 2D**). In the MeApd, *Slc32a*+ neurons exhibited greater *H1a*, but not *Arc*, expression response to a Dom over Sub (**Figure 2G**).

While stimulus and order/IEG channel interactions exhibited a lack of coherence across cell types, we did identify large-scale cell-type specific order effects throughout the amygdala. In the BLA, and MeApv, *Slc17a7+* neurons displayed significantly elevated IEG expression in response to P1 stimuli compared to P2 stimuli (P2/*Arc* – P1/*H1a* in BLA *Slc17a7+*:: β = −0.508 ± 0.025, p < 0.001; MeApv *Slc17a7+*: β = 0.4571 ± 0.115, p = 0.0003). This effect was also trending among EPd *Slc17a7+* neurons (P2/*Arc* – P1/H1a in EPd *Slc17a7+*: β = −0.079 ± 0.037, p = 0.078). In other words, *Slc17a7+* neurons exhibited more *H1a* expression than *Arc* expression, regardless of stimulus type. This effect was most pronounced among *Slc17a7+* BLA neurons. When we examined the paired responses to stimuli in each stimulus condition group, BLA *Slc17a7+* neurons exhibited a consistent decrease in IEG expression from P1 to P2 (See **Supplemental Figure S6 and Table S7** for within-subject IEG responses for each stimulus condition for all principal neuron types in each brain region). In contrast to the order effect observed among *Slc17a7+* neurons, we found that *Slc32a*+ neurons of the MeApd and MeAav exhibited an order effect in the opposite direction, such that *Arc* expression was greater than *H1a* expression (P2/*Arc* - P1/H1a in MeApd *Slc32a+* P2/*Arc* – P1/*H1a*: β = 0.090 ± 0.027, p = 0.002; P2/*Arc* – P1/H1a in MeAav *Slc32a+* P2/*Arc* - P1/*H1a:* β = 0.177 ± 0.076, p = 0.036).

### Cell Reactivation Patterns Indicate Recognition of Social Stimuli

We also determined whether overlapping or distinct ensembles exhibited IEG expression in response to pairs of stimuli. Cells re-activated by different social stimuli indicate general responsiveness to social information, while cells only re-activated by the same stimuli and not different stimuli, suggest a more specific detection and representation of distinct social cues.

Overlap, a measure of cell reactivation, was calculated for each brain region and cell population. We then applied GLMMs to test the association between overlap and each stimulus pairing and whether the pairing was social and concordant. Among *Slc17a7*+*Oxtr*+ EPd neurons, overlap trended higher for concordant social stimuli than discordant social stimuli (Social Discordant – Social Concordant: β = −0.069 ± 0.034, p = 0.05; Concordant – Discordant: β = −0.045 ± 0.034, p = 0.201) (**Figure 4A**). Among *Oxtr*+ DGh neurons, overlap was significantly greater for concordant stimuli than discordant stimuli (Social Concordant – Social Discordant: β =-0.143 ± 0.051, p = 0.010; Concordant – Discordant: β = −0.110 ± 0.048, p = 0.029) (**Figure 4B**). Approximately 12.6 ± 3.6% of DGh neurons were labeled for *Oxtr*. Notably, the opposite pattern was trending among *Slc17a7*+*Avrp1b*+*Oxtr*+ CA2CA3d neurons, where elevated overlap was observed among discordant social stimuli compared to concordant social stimuli (Social Concordant – Social Discordant: β =0.120 ± 0.058, p = 0.050; Concordant – Discordant: β = 0. 092 ± 0.052, p = 0.086) (**Figure 4C**) Approximately 17.8 ± 9.4% of *Slc17a7*+ CA2CA3d neurons were co-labeled for *Oxtr* and *Avrp1b*.

**Figure 4:**
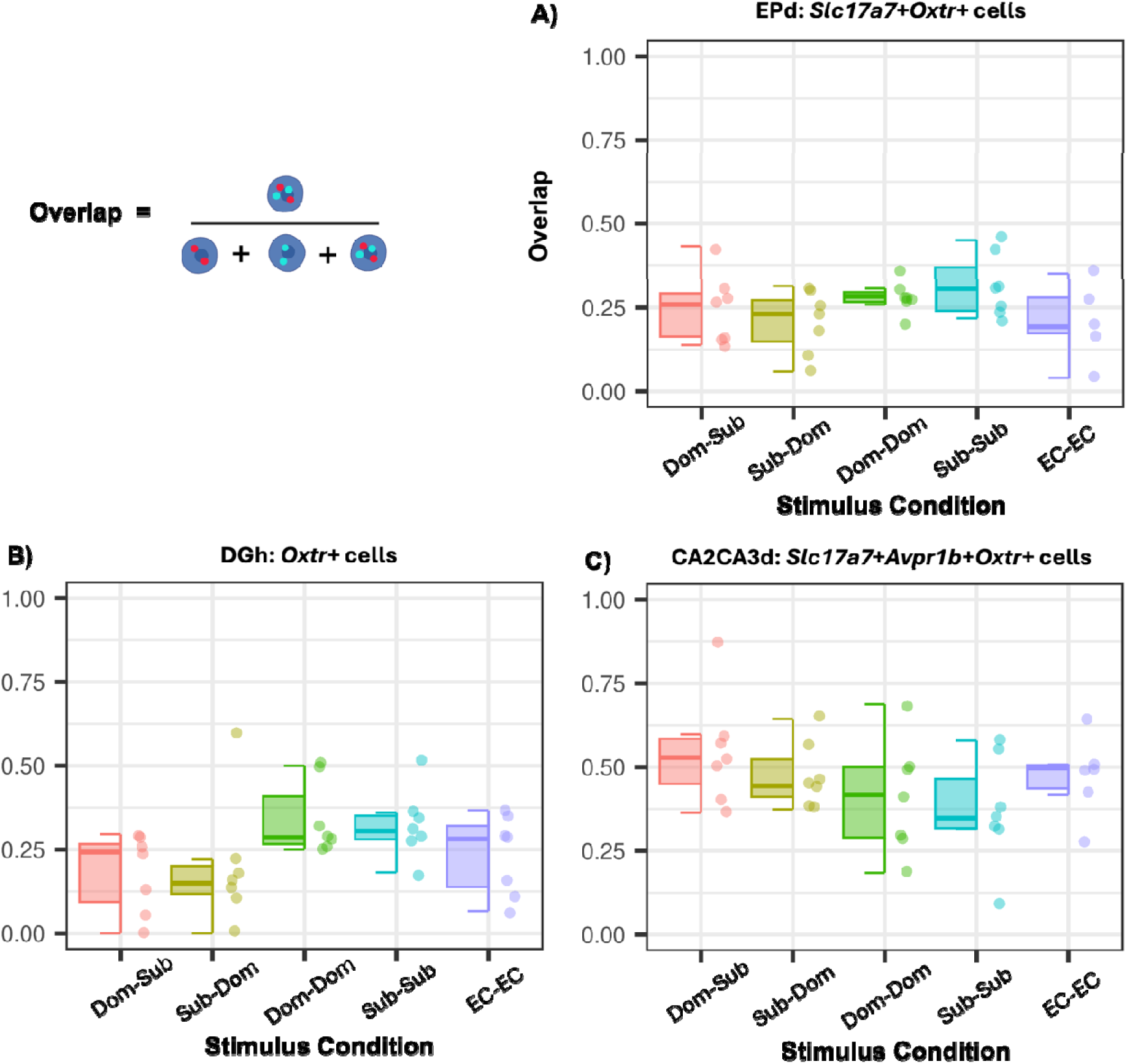
Cell Reactivation Across Stimulus Conditions. Overlap, a measure of cell reactivation, between ensembles of neurons activated by the first and second stimulus presentations. Significant associations between overlap and stimulus pairings were observed among *Slc17a7+* EPd neurons (**A**), *Oxtr*+ DGh neurons (**B**), and *Slc17a7+Avpr1b+Oxtr*+ CA2CA3 neurons (**C**).

## Discussion

We examined neural responses to dominant (Dom) and subordinate (Sub) stimuli to understand how the brain adapts to the dynamic social environment in dominance hierarchies. To achieve this, we used *Arc*/*H1a* catFISH to label neural ensembles activated by two social investigative events. We integrated this approach with RNAscope technology to label additional mRNA targets and characterize activated ensembles. We examined eight brain regions spanning the amygdala and hippocampus: BLA, MeApd, MeAav, MeApv, EPd, CA2CA3d, CA3p, and the DGh. These regions have been implicated in various functions necessary for social processing, including olfactory perception, emotional regulation, and social recognition (Raam and Hong 2021; Raam et al. 2017; Wang et al. 2014).

Throughout the amygdala, we observed elevated responses to Dom compared to Sub and the non-social empty cage control (EC). The responses of all *Slc17a7*+ BLA neurons were linearly ordered, with greater responses to Dom than to Sub and EC, and greater responses to Sub than to EC. We observed this same pattern among *Slca17a7*+*Drd1*+ neurons. It has been proposed that BLA-centric mechanisms underlying fear learning contribute to learning social ranks within dominance hierarchies (Dwortz et al. 2022). Supporting evidence stems from studies in macaques that showed that the firing rate of BLA neurons correlates with the social ranks of familiar hierarchy members’ faces (Munuera, Rigotti, and Salzman 2018). While the role of dopamine in the BLA has not been specifically studied in the context of dominance hierarchies and social learning, its multifaceted role in Pavlovian fear learning is well-documented (Heath et al. 2015; Fadok et al. 2010). This role includes driving attention during learning by enhancing the salience of stimuli (Nieoullon 2002). In dominance hierarchies, attention is often directed toward more dominant group members (Chance and Larson 1976, Freniere and Charlesworth 1983; Vaughn and Waters 1981; Pannozzo et al. 2007; Curley 2016). This monitoring of dominant individuals facilitates learning from successful individuals and threat avoidance. Thus, elevated responses of *Slca17a7*+ BLA neurons, and *Slca17a7*+*Drd1*+ neurons specifically, to Dom may also reflect attentional processes and the salience of dominant individuals.

We also observed response patterns similar to the BLA in the EPd, a relatively understudied brain region bordering the BLA. The EPd is highly heterogeneous, containing glutamatergic and GABAergic cells and oxytocin and D1 and D2 dopamine receptors (Dubois et al. 1986; Savasta, Dubois, and Scatton 1986; Biggs and Hammock 2022). We found that glutamatergic *Slc17a7*+ EPd neurons exhibited the most IEG expression in response to Dom, followed by Sub, and lastly, EC. We found that many of these neurons were co-labeled for *Oxtr*. Consistent with these findings, a study in juvenile mice found high colocalization of *Oxtr* and *Slc17a7* mRNA in the EP, and that application of oxytocin on EP cells stimulates glutamatergic activity (Biggs and Hammock 2022). The precise function of the EP has yet to be thoroughly investigated. It likely serves numerous functions given its many reciprocal connections with areas involved in olfaction (e.g., olfactory bulb, medial amygdala, perirhinal cortex), emotional responses (BLA), memory (e.g., entorhinal cortex), and even sensorimotor functions (Neafsey, Hurley-Gius, and Arvanitis 1986; Behan and Haberly 1999; Meis et al. 2008; Watson, Smith, and Alloway 2017). There is also evidence to suggest that *Oxtr*+ EPd neurons may facilitate social processing. For example, adult mice exhibit greater OxtR binding in the EP after communal rearing compared to being raised by a single mother, suggesting that EP OxtR is upregulated in contexts where social processing load is more demanding (Curley et al. 2009).

Elevated responses to dominant stimuli were observed in GABAergic *Slc32a*+ neurons in the MeAav. This pattern persisted in the MeApv, where *Slc32a*+ innervation was more scArce. Notably, we did not observe stimulus-associated responses among glutamatergic *Slc17a7*+ MeApv neurons. Thus, socially receptive neurons in the highly heterogeneous MeA appear to be primarily GABA-ergic. This observation extends our previous finding that the MeApv exhibits elevated cFos immunoreactivity in response to dominant olfactory cues (Lee et al. 2021). Numerous studies have also shown that GABAergic neurons are the primary population of MeA projection neurons that drive social responses, including aggression (Li et al. 2017; Choi et al. 2005; Hong, Kim, and Anderson 2014; Keshavarzi et al. 2014; Raam and Hong 2021). Thus, GABA-ergic MeA neurons may mediate defensive responses to more dominant hierarchy members. While subjects did not fight during stimulus presentations, response patterns may reflect social decision-making and the initiation of such behavior under the appropriate social context. Furthermore, we found that *Slc32a+Oxtr*+ neurons in the MeApd and MeAav exhibited an elevated response to Dom. We did not observe this pattern for all unique subpopulations (e.g., *Slc32a+Drd1*+ neurons). It has been previously demonstrated that oxytocin action in the MeApd mediates the formation of representations of sociosexual olfactory cues (Li et al. 2017). Our findings suggest that *Slc32a+Oxtr*+ MeA neurons may play a similar role in representations of dominance cues.

We also assessed the effect of stimulus presentation duration on IEG expression. Stimulus presentations varied in duration due to stimulus animals being removed early to avoid fighting. Responses of *Slc17a7+* neurons in CA2CA3d and CA3p, *Slc17a7+* BLA neurons, and *Slc32a+* MeApd and MeAav neurons were positively associated with presentation duration. Notably, presentation duration was not associated with the social status of the stimulus animal. One simple explanation for this finding is that the number of cells activated is directly proportional to the amount of sensory input. However, a second possibility is that these populations are ostensibly time-locked to the end of stimulus presentations. One study has found that activity in the dorsal hippocampus and caudate nucleus is time-locked to stimulus offset reflecting post-stimulus processes important for memory encoding (Ben-Yakov and Dudai 2011). In the present study, *H1a* and *Arc* transcription were halted at 30 minutes and 5 minutes from the onset of P1 and P2, respectively. While there was no individual variation in the timing of the stimulus onset, there was variation in the offset, as animals were removed to avoid fighting. Thus, subjects sacrificed closer to the end of their stimulus presentations exhibited higher levels of IEG expression.

We also identified prominent order effects in the amygdala, in which one order/IEG channel was elevated compared to the other. In MeApd and MeAav *Slc32a+* neurons, the expression of *Arc* was greater than *H1a*. Conversely, in BLA and MeApv *Slc17a7*+ neurons, expression of *H1a* was greater than *Arc*. While P1 durations trended longer than those for P2, this likely does not explain order effects. If this were the case, we would also expect to find elevated *H1a* expression in *Slc32a+* MeApd and MeAav populations, which also exhibited responses positively associated with presentation duration. Alternatively, these order effects could be relevant to subjects’ perception of stimuli. Reduced *Arc* among *Slc17a7*+ neurons in the amygdala could reflect habituation or the subjects’ reduced attention to or perceived salience of stimuli in P2. A third and perhaps most likely explanation is that these order effects could be due to the intrinsic properties of these cell types. H1a has been identified in GABA-ergic populations, but its function at inhibitory synapses remains less understood than its role at glutamatergic synapses (Xiao et al. 1998; Imamura et al. 2011). Thus, *H1a* may be downregulated compared to *Arc* in inhibitory neurons.

We also observed interactions between stimulus-evoked responses and presentation order/IEG channels, such that stimulus-associated response patterns were more robust with respect to P1/H1a or P2/Arc. For example, among *Slc32a+* MeApd neurons, *H1a*, but not *Arc*, expression was significantly elevated in response to a dominant stimulus over a subordinate and empty cage. Unlike the general order effects, interactions between order and social context did not exhibit a coherent association with cell-type. These observed order effects and interactions provide valuable insights and methodological considerations for IEG-based studies on social processing in the brain. Multiple factors shape IEG expression, including the time course of stimuli and the cell type-specific properties that may lead certain IEGs to be better indicators of activity. Hence, methods that allow reseArchers to capture multiple IEG responses provide a more thorough picture of how the brain responds to stimuli. Such methods include catFISH as well as the latest advancements in single-cell RNA sequencing, which allows reseArchers to explore numerous IEGs and select optimal candidates for various cell types (Wu et al. 2017).

In addition to evaluating levels of IEG expression across stimulus types, we also examined overlap, or the reactivation of neural ensembles across stimulus pairings. *Slc17a7*+*Oxtr*+ EPd neurons exhibited elevated overlap among concordant social stimulus pairs (i.e. Dom-Dom and Sub-Sub) compared to discordant social stimulus pairs (Dom-Sub and Sub-Dom). Taken together with the finding that *Slc17a7*+*Oxtr*+ EPd neurons respond more to dominant stimuli, it appears that these cells’ activity could represent the salience of social context. The EPd is known to be generally involved in olfactory processing, and our findings further suggest that it serves an important role in processing social olfactory cues specifically. Comprised of diverse neuropeptide receptor-expressing cells and connections with wide-spread brain regions, from olfactory nuclei to sites of emotional processing, the EPd is a region worth further attention in future social olfaction studies.

We also identified significant reactivation patterns in hippocampal subregions that may shed light on memory processes engaged during social investigation of dominance hierarchy members. Specifically, we observed that overlap among *Oxtr*+ neurons in the DGh was higher for concordant stimuli compared to discordant stimuli. This aligns with the DGh’s role in pattern separation—a process through which distinct inputs from the entorhinal cortex are transformed into unique representations, facilitating the encoding and recollection of specific social stimuli (McHugh et al., 2007; Raam et al., 2017). The presence of *Oxtr*+ neurons in the DGh further supports this role, as previous work has shown that the deletion of *Oxtr* in DGh mossy cells disrupts social recognition, highlighting its necessity in social memory formation (Raam et al., 2017, Hung et al., 2023).

In a previous catFISH experiment, Raam et al. identified overlap patterns in CA3p that resemble what we observed in the DGh. They showed that mice presented with the same familiar animal twice exhibit a higher degree of overlap among CA3p neurons than those presented with a familiar and novel individual (Raam et al. 2017). This finding supported the knowledge that CA3p neurons perform pattern separation on DGh inputs to distinguish between different social stimuli and facilitate individual recognition. It has also been shown that CA3 cells perform attractor dynamics on DG inputs and pattern completion when stimuli are the same or only slightly different to facilitate memory retrieval (McHugh et al. 2007; Guzowski, Knierim, and Moser 2004). One explanation for why we did not observe higher overlap in CA3 in response to two identical stimuli is that two familiar inputs from the same social group give rise to pattern completion and generalized representations that could reflect the recognition of familiar group members. It is also possible that separate dominant and subordinate representations would emerge over a longer investigative period or with methods that provide better temporal precision (e.g., electrophysiology). Additionally, activity in the DGh is sparse compared to activity in CA3p and CA2CA3d, and perhaps this resulted in clearer stimulus-evoked *H1a* and *Arc* expression.

Interestingly, we observed an inverse pattern in CA2CA3d compared to DGh, with *Slc17a7+Avrp1b+Oxtr*+ neurons showing higher overlap for discordant stimuli. Notably, *Avrp1b* is a marker for the CA2 subregion, which is known to support social memory in rodents (Hitti and Siegelbaum 2014), and our findings confirmed abundant *Avrp1b* labeling in this region (see **Supplemental Figure S7**). Given CA3’s ability to flexibly perform pattern separation or pattern completion based on input similarity, this pattern may reflect attractor dynamics facilitating pattern completion, resulting in the absence of ensemble distinction for distinct stimuli beyond the DGh. Taken together, the activity patterns we observed in the DGh and CA2CA3 regions may be indicative of encoding social memories of specific hierarchy members. It is now a well-substantiated theory that memory formation involves the aggregation of neurons into distinct ensembles, or “engrams,” that represent particular memories and are reactivated during recall (McHugh et al. 2007; Raam et al. 2017). Engrams associated with various types of memories, including individual social recognition, have been well-documented in the rodent hippocampus, specifically in ventral CA1 (Okuyama 2018). Evidence suggests that these distinct ensemble activity patterns are shaped by inputs from the dorsal hippocampus (Watarai et al. 2021). Collectively, these findings support the role of dorsal hippocampal regions in processing and encoding social information within the context of dominance hierarchies.

## Conclusion

We identify neural populations involved in processing social information in dominance hierarchies. The amygdala and Epd exhibited robust responses to social stimuli, with distinct subpopulations showing heightened sensitivity to dominant stimuli. *Slc17a7+* BLA neurons, which exhibited co-labeling of *Drd1*+, *Slc32a+* MeA populations with abundant *Oxtr+* labeling and *Slc17a7+* EPd neurons co-labeled for *Oxtr*+ and *Drd1*+ displayed the highest levels of IEG expression in response to a dominant stimulus. The catFISH approach allowed us to capture these shifts in IEG expression between and within subjects. It also enabled us to examine the reactivation of ensembles, between stimulus types, revealing potential sites of engrams for familiar hierarchy members. We found that distinct *Slc17a7+Oxtr+* ensembles in the EPd respond to dominant and subordinate individuals, while more similar, or overlapping ensembles respond to the same individual. In the hippocampus, we found evidence of memory storage processes that may facilitate recognition of dominance hierarchy members. Specifically, *Oxtr*+ DGh ensembles exhibit less reactivation in subjects presented with dominant and subordinate individuals, while the inverse pattern was observed among *Slc17a7*+ *Avrp1b*+*Oxtr*+ CA2CA3d neurons. Overall, our findings detail how the amygdala and hippocampus process social information at the level of distinct cell populations and shed light on these regions’ role in adapting to dynamic social environments. Finally, in this study we also demonstrate the effective integration of catFISH and RNAscope technology and provide considerations for future applications of this approach. These considerations include the impact of stimulus onset and offset on IEG transcriptional timelines, as well as the relevance of cell-type characteristics in determining candidate IEGs.

## Acknowledgements

The authors wish to acknowledge funding from NIH grant 5R21MH132981-02.

## Data and Code Accessibility

All data, code and analyses are publicly available at https://github.com/mfdwortz/catFISH

---

**Figure S1:**
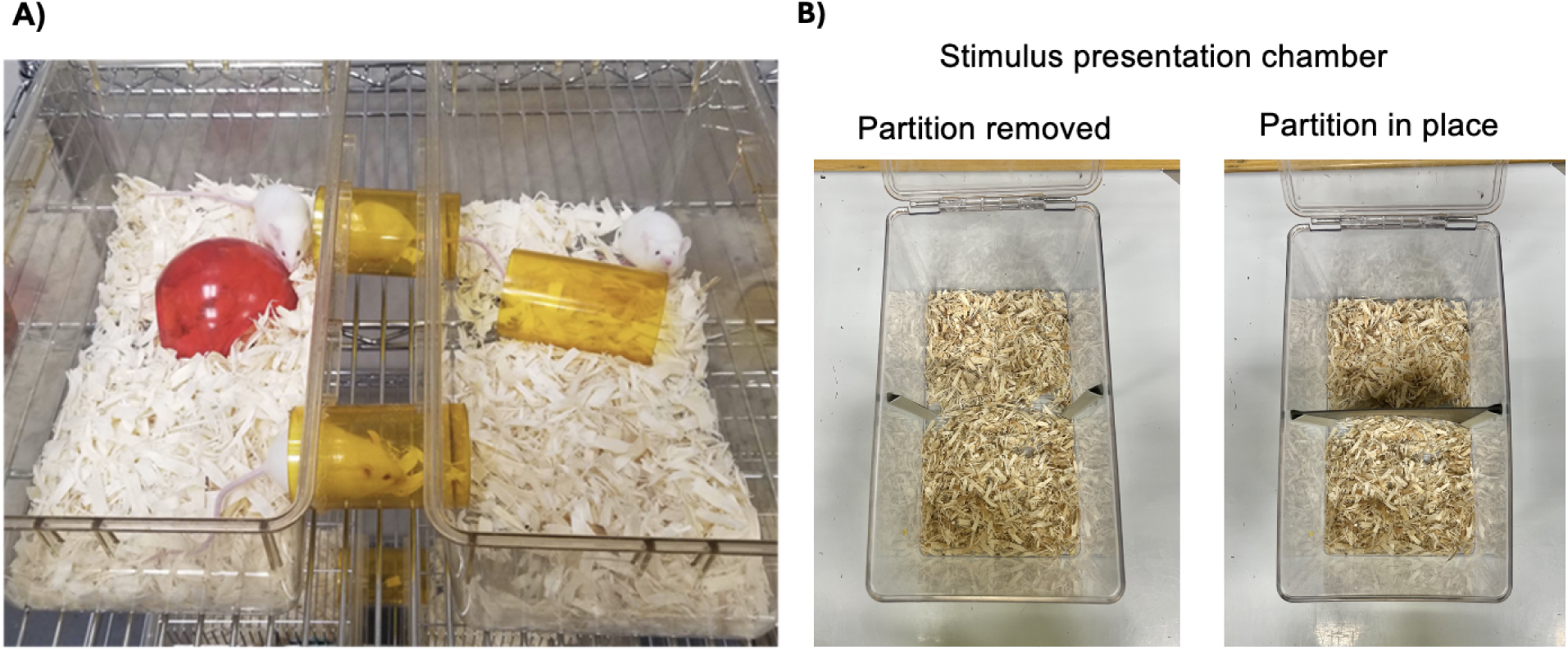
Social Housing and Stimulus Presentation Chambers. **A**) Image of group housing structure consisting of two standard-size rat cages connected by acrylic tubes. **B**) Top-down view of stimulus presentation chamber with and without the partition.

**Figure S2:**
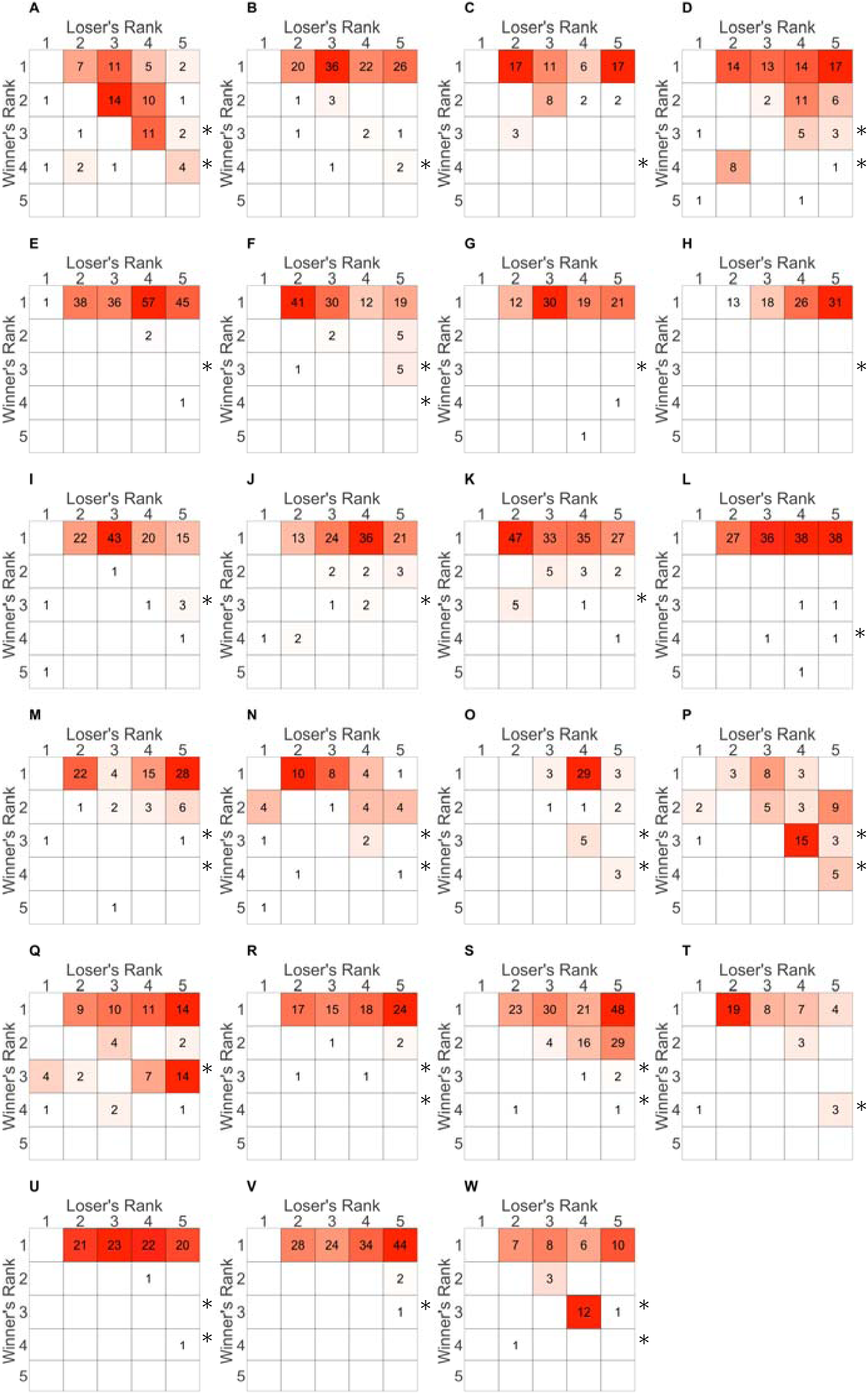
Sociomatrices. Sociomatrices show the total frequency of agonistic interactions that occurred between all pairs of individuals across all cohorts (A-J) over the entire observation period. Winners of each contest are listed in rows, and losers are listed in columns. Ranks were calculated using the DS method (see methods). Cells of each matrix are colored on a gradient from white (lowest value in each matrix) to red (highest value in each matrix).

**Figure S3:**
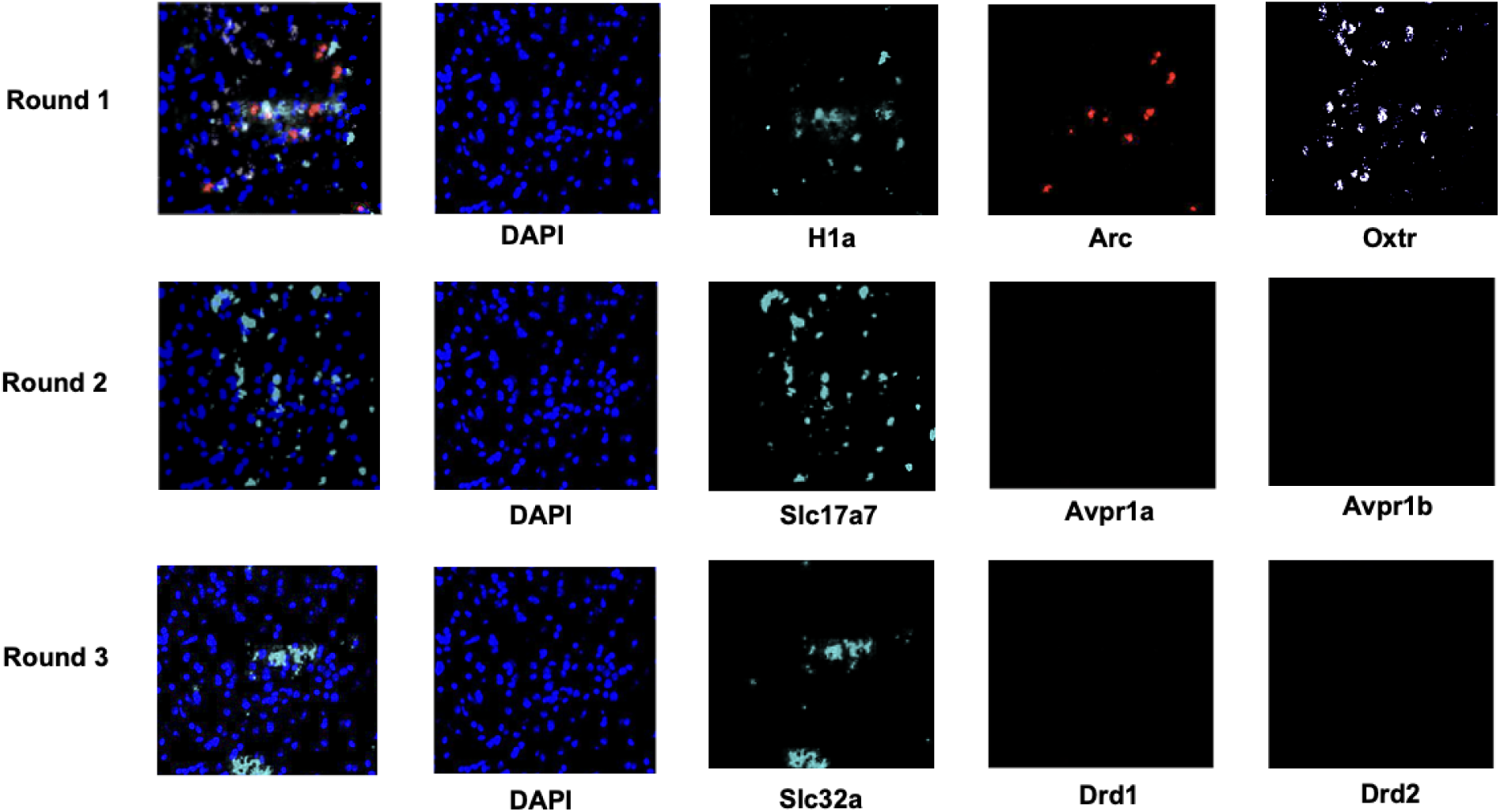
RNAscope Image Registration. RNAscope labeling involves three rounds of labeling and imaging, during which fluorophores are sequentially cleaved. Images for each mRNA target are aligned using DAPI staining from each round and merged into a composite image.

**Figure S4:**
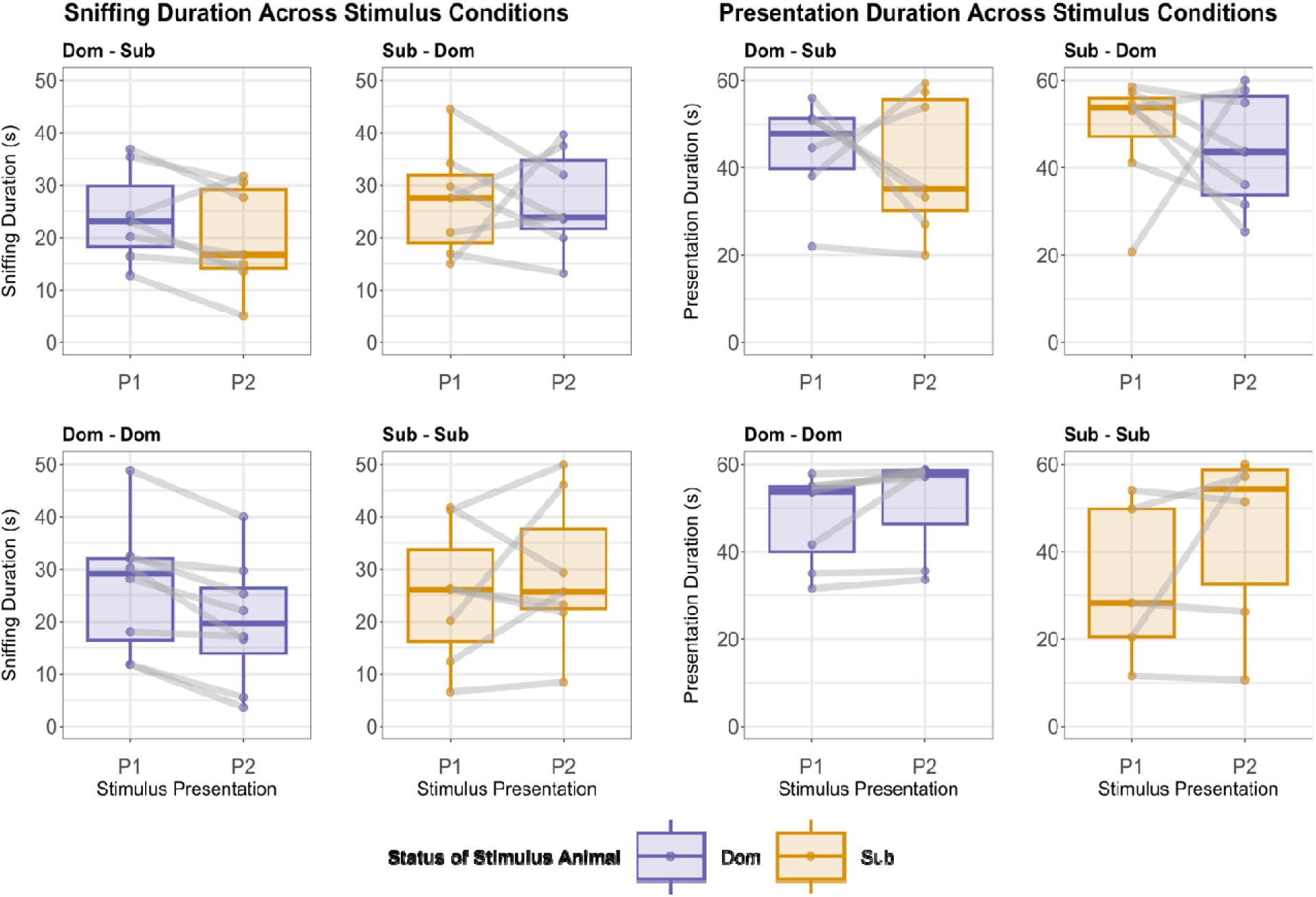
Sniffing and Presentation Duration in Each Stimulus Condition. Sniffing and presentation duration during the first (P1) and second (P2) stimulus presentation in each stimulus condition where animals were presented with a pair of hierarchy members. Significance levels for each comparison are shown (p < 0.05 (*), p < 0.01 (**), p < 0.001 (***).

**Figure S5:**
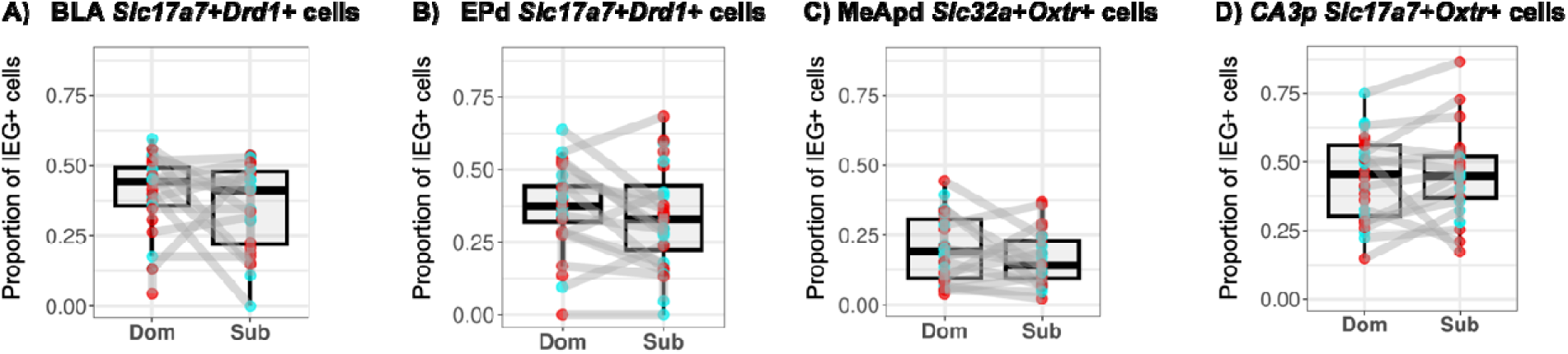
Within-Subject Differences in IEG Expression Between Dom and Sub Stimuli. Boxplots illustrate within-subject differences in IEG expression levels between dominant (Dom) and subordinate (Sub) stimuli across various cell populations. In these populations, differences in IEG expression between Dom and Sub stimuli do not appear visually significant (e.g., BLA *Slc17a7*+*Drd1*+ cells; see Figure 2C). Subject ID was included as a random effect in the statistical model, allowing within-subject changes in IEG expression between stimuli to contribute to significant differences overall.

**Figure S6:**
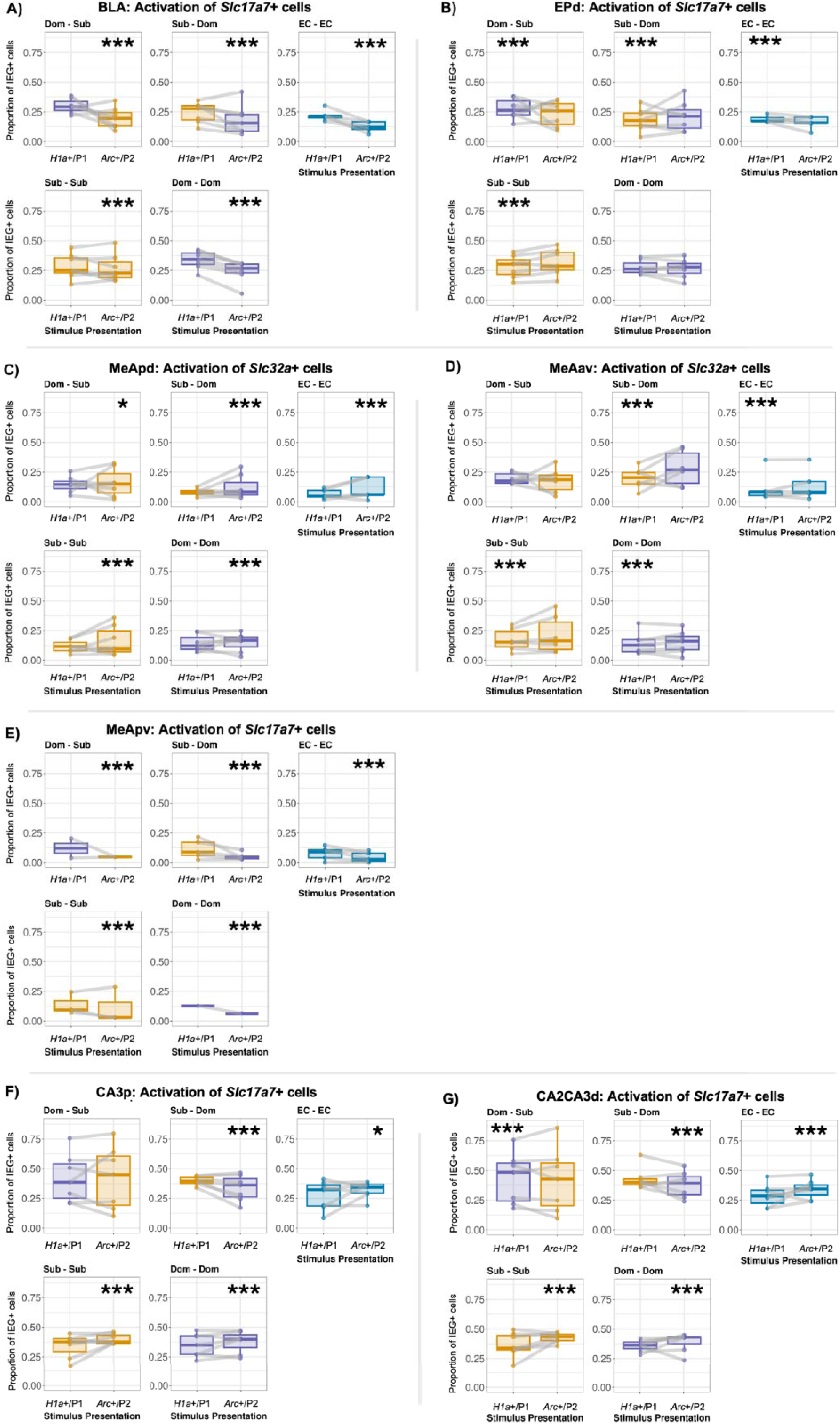
Within-Subject IEG Responses in Each Stimulus Condition. Boxplots show IEG responses for principal neurons of the BLA (A), EPd (B), MeApd (C), MeAav (D), MeApv (E), CA3p (F), and CA2CA3d (G). Binomial GLMMs were applied to test the association between the proportion of IEG-expressing cells and the stimulus presentation order (P1 vs P2), and the significance level is shown (p < 0.05 (*), p < 0.01 (**), p < 0.001 (***).

**Figure S7:**
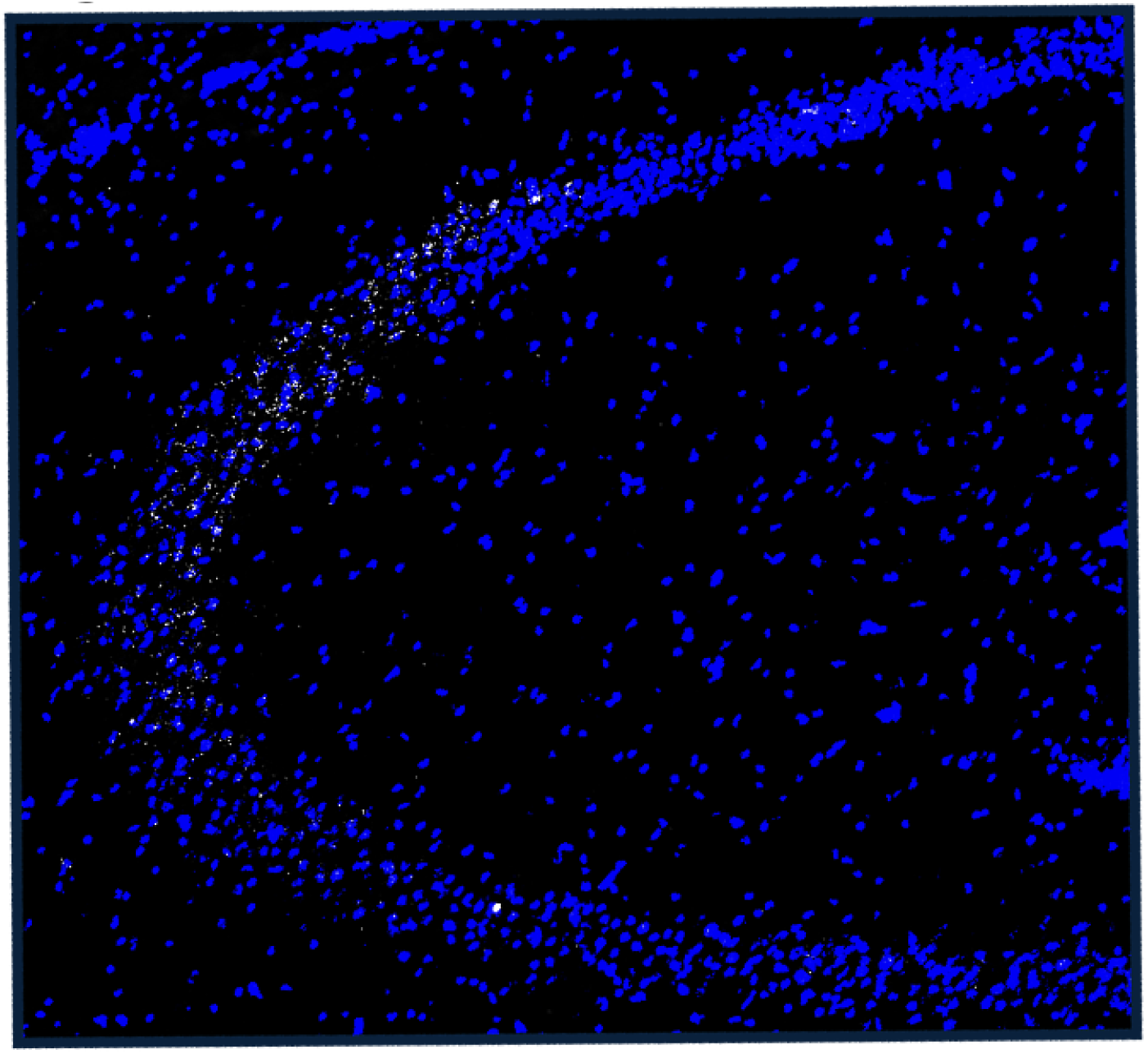
Elevated *Avpr1b* Labeling in CA2. RNAscope labeling of *Avpr1b* is shown in white, highlighting increased expression in CA2. Blue DAPI staining indicates nuclei.

**Table S1:** Mouse Social Behavior Ethogram. Observers documented instances of agonistic interactions by recording pairs of offensive and defensive behaviors (e.g., “Bite/Flight - Flee”). In cases of a “pre-emptive flee,” where the aggressive animal displayed no observable behavior, “NA” was recorded.

| Dominant Behaviors | Description | Subordinate Behaviors | Description |
| --- | --- | --- | --- |
| **Bite/Fight** | Individual initiates biting and wrestling. | **Subordinate posture** | Individual sits upright or rolls over and raised forepaws, exposing the abdomen. |
| **Lunge** | Individual rapidly jumps towards another. | **Flee** | Individual quickly moves away from another performing an aggressive behavior. |
| **Chase** | Individual rapidly follows a fleeing target. | **Pre-emptive flee** | Individual quickly moves away from another animal who has not exhibited threatening behavior. |
| **Mount** | Individual mounts another in the absence of intromission or breeding opportunity. | **Freeze/immobility** | Individual ceases movement and does not flee. Freezing can be indiscernible from immobility when the individual is restrained by an aggressor. |

**Table S2:** Model Selection Results. . Models a and b were applied to all brain regions and cell populations therein. We also applied more complex models (c through l) to evaluate possible influences of presentation order and stimulus durations on IEG expression.

| Candidate models |  |  |
| --- | --- | --- |
| Model id | Formula |  |
| a | social + (1 \| Subject ID) + (1 \| order) + (1 \| batch) |  |
| b | stimulus type + (1 \| Subject ID) + (1 \| order) + (1 \| batch) |  |
| c | social + order + (1 \| Subject ID) + (1 \| batch) |  |
| d | stimulus type + order + (1 \| Subject ID) + (1 \| batch) |  |
| e | social * order + (1 \| Subject ID) + (1 \| batch) |  |
| f | stimulus type * order + (1 \| Subject ID) + (1 \| batch) |  |
| g | stimulus type + presentation duration + (1 \| Subject ID) + (1 \| order) + (1 \| batch) |  |
| h | stimulus type + sniff duration + (1 \| Subject ID) + (1 \| order) + (1 \| batch) |  |
| i | stimulus type * presentation duration + (1 \| Subject ID) + (1 \| order) + (1 \| batch) |  |
| j | stimulus type * sniff duration + (1 \| Subject ID) + (1 \| order) + (1 \| batch) |  |
| k | (stimulus type + presentation duration) * order + (1 \| Subject ID) + (1 \| batch) |  |
| l | (stimulus type + sniff duration) * order + (1 \| Subject ID) + (1 \| batch) |  |
| Optimal models |  |  |
| Brain region | Cell type(s) | Optimal model(s) |
| BLA | all cells+\| oxtr+\| Slc32a+\| Slc17a7+\| Slc17a7,drd1+ | f, k |
| BLA | drd1+ | f, i |
| BLA | oxtr,Slc17a7+ | e, k |
| Ca2Ca3d | all cells+\| Slc17a7+ | f, i |
| Ca2Ca3d | oxtr+\| oxtr,Slc17a7+ | f, k |
| Ca2Ca3d | oxtr,Slc17a7,avpr1b+\| Slc17a7,avpr1b+\| avpr1b+ | c, k |
| Ca3p | all cells+\| Slc17a7+ | f, i |
| Ca3p | oxtr,Slc17a7+\| oxtr+ | d, k |
| Ca3p | Slc17a7,avpr1b+\| avpr1b+ | e, l |
| DGhilus | all cells+\| oxtr+ | e, k |
| DGhilus | oxtr,Slc32a+\| Slc32a+ | e |
| DGhilus | Slc17a7+ | e, l |
| EPd | all cells+\| oxtr,Slc17a7+\| Slc17a7+ | f, k |
| EPd | oxtr+\| Slc32a+\| Slc17a7,drd1+ | d, k |
| EPd | oxtr,drd1+\| drd1+ | d, l |
| EPd | oxtr,Slc17a7,drd1+ | d, j |
| EPd | oxtr,Slc17a7,drd2+ | c |
| EPd | drd2+ | e, k |
| MeAav | all cells+\| oxtr,Slc32a+\| Slc32a+ | f, k |
| MeAav | drd1+ | f, j |
| MeAav | oxtr+ | f, l |
| MeAav | Slc32a,drd1+ | d, l |
| MeApd | all cells+\| oxtr+\| oxtr,Slc32a+\| Slc32a+ | f, k |
| MeApv | all cells+ | f, l |
| MeApv | oxtr+ | e, l |
| MeApv | oxtr,Slc32a+ | d, j |
| MeApv | oxtr,Slc17a7+\| Slc17a7+ | e, i |
| MeApv | Slc32a+ | f, j |

**Table S3:** Dominance Results. Directional Consistency (DC) of dominance interactions and David’s Scores (DS) were calculated. In brief, DC assesses the degree to which all agonistic interactions in a group occur in the direction of the more dominant individual to the more subordinate individual within each relationship. It is equal to (H-L)/(H+L) where H is the frequency of behaviors occurring in the most frequent direction and L is the frequency of behaviors occurring in the least frequent direction within each relationship. The significance of DC values were evaluated using the randomization test proposed by Leiva, Solanas, & Salafranca (2008). DS provides an individual dominance rating and ranking for each individual in a group, determining the overall success of each individual at winning contests relative to the success of all other individuals. The most dominant and subordinate individuals in each hierarchy had the highest and lowest DS, respectively. These individuals served as stimulus animals. Subjects were selected from mid-ranked individuals with the third and fourth-highest DS.

| Cohort | DC | Rank 1 ID | Rank 2 ID | Rank 3 ID | Rank 4 ID | Rank 5 ID | Rank 1 DS | Rank 2 DS | Rank 3 DS | Rank 4 DS | Rank 5 DS |
| --- | --- | --- | --- | --- | --- | --- | --- | --- | --- | --- | --- |
| A | 0.836 | 2 | 5 | 1 | 3 | 4 | 7.054 | 3.153 | -0.733 | -2.89 | -6.583 |
| B | 0.965 | 4 | 5 | 1 | 3 | 2 | 7.7 | -0.046 | -1.407 | -1.532 | -4.715 |
| C | 0.909 | 5 | 1 | 4 | 3 | 2 | 6.438 | 0.264 | -1.766 | -2.337 | -2.599 |
| D | 0.773 | 4 | 5 | 3 | 1 | 2 | 8.772 | 1.851 | 0.292 | -4.792 | -6.123 |
| E | 1 | 3 | 5 | 4 | 1 | 2 | 5.867 | 0.123 | -0.965 | -2.22 | -2.805 |
| F | 0.983 | 3 | 4 | 5 | 2 | 1 | 6.711 | 0.188 | -0.791 | -0.861 | -5.247 |
| G | 0.976 | 1 | 2 | 5 | 4 | 3 | 4.748 | -0.872 | -1.006 | -1.43 | -1.441 |
| H | 1 | 3 | 2 | 4 | 5 | 1 | 3.808 | -0.882 | -0.938 | -0.985 | -1.002 |
| I | 0.963 | 4 | 2 | 1 | 3 | 5 | 7.32 | 0.372 | -0.583 | -2.403 | -4.706 |
| J | 0.943 | 1 | 2 | 5 | 3 | 4 | 7.515 | 0.137 | -1.881 | -2.784 | -2.988 |
| K | 0.937 | 3 | 1 | 4 | 5 | 2 | 8.748 | 0.76 | -0.111 | -4.306 | -5.092 |
| L | 0.972 | 3 | 4 | 1 | 5 | 2 | 6.808 | -0.95 | -0.95 | -1.954 | -2.954 |
| M | 0.952 | 2 | 1 | 3 | 4 | 5 | 6.729 | 1.781 | -2.073 | -2.724 | -3.713 |
| N | 0.667 | 5 | 2 | 3 | 4 | 1 | 4.4 | 2.85 | -0.683 | -2.983 | -3.583 |
| O | 1 | 3 | 5 | 1 | 2 | 4 | 4.583 | 2.35 | 0.667 | -3.1 | -4.5 |
| P | 0.895 | 1 | 5 | 2 | 4 | 3 | 5.425 | 5.283 | 0.36 | -4.052 | -7.017 |
| Q | 0.778 | 3 | 5 | 1 | 2 | 4 | 6.672 | 1.762 | 0.516 | -1.949 | -7 |
| R | 0.975 | 1 | 4 | 2 | 5 | 3 | 6.625 | -0.637 | -0.78 | -2.416 | -2.791 |
| S | 0.989 | 2 | 4 | 5 | 3 | 1 | 9.651 | 4.104 | -1.503 | -4.47 | -7.782 |
| T | 0.956 | 3 | 4 | 1 | 5 | 2 | 4.847 | 0.344 | -1.014 | -1.222 | -2.956 |
| U | 1 | 4 | 1 | 5 | 2 | 3 | 5.732 | -0.181 | -0.964 | -1.912 | -2.675 |
| V | 1 | 1 | 4 | 3 | 2 | 5 | 5.815 | -0.365 | -0.765 | -0.977 | -3.709 |
| W | 1 | 5 | 2 | 3 | 4 | 1 | 7.05 | -0.608 | -1.126 | -2.57 | -2.746 |

**Table S4:** Evaluating Effect of Order and Status of Stimulus Animal on Presentation and Sniffing Duration. Applied models and results testing the effect of stimulus presentation order (P1 vs. P2) and status of stimulus animals (Dom vs. Sub) on the duration of presentations and the duration of sniffing.

| Model: Presentation Duration ~ Status * Order + (1\|Subject ID) |  |  |  |  |  |  |  |
| --- | --- | --- | --- | --- | --- | --- | --- |
| Fixed effect | Estimate | Std. Error | df | t value | Pr(>\|t\|) | 0.025 | 0.975 |
| stim_status | -1.398 | 5.075 | 45.845 | -0.275 | 0.783 | -11.189 | 8.354 |
| stim_status:stimulus_order | -2.012 | 7.128 | 44.914 | -0.282 | 0.779 | -15.764 | 11.683 |
| stimulus_order | 1.307 | 4.579 | 36.453 | 0.285 | 0.775 | -7.499 | 10.171 |
| Model: Sniffing Duration ~ Status * Order + (1\|Subject ID) |  |  |  |  |  |  |  |
| Fixed effect | Estimate | Std. Error | df | t value | Pr(>\|t\|) | 0.025 | 0.975 |
| stimulus_status | -4.795 | 3.269 | 36.238 | -1.467 | 0.142 | -11.197 | 1.828 |
| stimulus_status:stimulus_order | 7.496 | 4.572 | 35.563 | 1.639 | 0.11 | -1.692 | 16.398 |
| stimulus_order | -5.388 | 2.845 | 31.257 | -1.894 | 0.058 | -10.872 | 0.278 |
| Model: Sniffing Duration ~ Location of Stimulus Animal + Stimulus Status + Order + (1\|Subject ID) |  |  |  |  |  |  |  |
| Fixed effect | Estimate | Std. Error | df | t value | Pr(>\|t\|) | 0.025 | 0.975 |
| sniff_back | 8.917 | 1.183 | 141.215 | 7.535 | 0 | 6.612 | 11.221 |
| sniff_front | -1.803 | 1.212 | 143.305 | -1.487 | 0.139 | -4.19 | 0.55 |
| stim_type | 0.387 | 1.161 | 137.687 | 0.333 | 0.74 | -1.926 | 2.695 |
| stimulus_order | -0.555 | 0.979 | 141.002 | -0.568 | 0.571 | -2.461 | 1.35 |
| Post-hoc |  |  |  |  |  |  |  |
| Comparison |  | Estimate | Std. Error | t value | Pr(>\|t\|) |  |  |
| sniff_back - sniff body |  | 8.917 | 1.183 | 7.535 | 0 |  |  |
| sniff_front - sniff body |  | -1.803 | 1.212 | -1.487 | 0.137 |  |  |
| sniff_front - sniff_back |  | -10.72 | 1.219 | -8.791 | 0 |  |  |
| Model: P2 Duration ~ P1 Stimulus Status |  |  |  |  |  |  |  |
|  | Fixed effect | Estimate | Std. Error | t value | Pr(>\|t\|) | 0.025 | 0.975 |
| Dom stimulus animal in P2 | P1_stimulus_status | -8.941 | 6.552 | -1.365 | 0.196 | -23.096 | 5.215 |
| Sub stimulus animal in P2 | P1_stimulus_status | 5.896 | 9.672 | 0.61 | 0.554 | -15.178 | 26.97 |
| Model: P2 Sniff Duration ~ P1 Stimulus Status |  |  |  |  |  |  |  |
|  | Fixed effect | Estimate | Std. Error | t value | Pr(>\|t\|) | 0.025 | 0.975 |
| Dom stimulus animal in P2 | P1_stimulus_status | 7.035 | 5.701 | 1.234 | 0.239 | -5.281 | 19.351 |
| Sub stimulus animal in P2 | P1_stim_type | 9.185 | 6.648 | 1.382 | 0.192 | -5.301 | 23.67 |

**Table S5:** Comparison of Presentation and Sniffing Duration within Each Stimulus Condition. Applied models and results testing the effect of stimulus presentation order (P1 vs. P2) and status of stimulus animals (Dom vs. Sub) on the duration of presentations and sniffing within each social stimulus presentation condition.

| Model: Presentation Duration ~ Order + (1\|Subject ID) |  |  |  |  |  |
| --- | --- | --- | --- | --- | --- |
| Fixed effect | Comparison | Estimate | Std. Error | t value | Pr(>\|t\|) |
| Dom-Sub | **P2 - P1** | -5.4003 | 6.4783 | -0.8336 | 0.4045 |
| Sub-Dom | **P2 - P1** | -4.288 | 7.2606 | -0.5906 | 0.5548 |
| Dom-Dom | **P2 - P1** | 5.163 | 2.1007 | 2.4577 | 0.014 |
| Sub-Sub | **P2 - P1** | 5.1766 | 5.7529 | 0.8998 | 0.3682 |
| Model: Sniffing Duration ~ Order + (1\|Subject ID) |  |  |  |  |  |
| Dom-Sub | **P2 - P1** | -4.1181 | 2.2403 | -1.8382 | 0.066 |
| Sub-Dom | **P2 - P1** | 0.0716 | 5.1017 | 0.014 | 0.9888 |
| Dom-Dom | **P2 - P1** | -6.6279 | 1.392 | -4.7613 | <0.001 |
| Sub-Sub | **P2 - P1** | 4.2913 | 4.8411 | 0.8864 | 0.3754 |

**Table S6:** Stimulus Presentation Order Effects Shape Responses to Social Stimuli. Statistical results from the following lmer model are shown: stimulus type * order + (1 | Subject ID) + (1 | batch). We tested the interaction between stimulus type and order. Order refers to the IEG/Presentation order channel (e.g. h1a/P1 and arc/P2). Post-hoc comparisons were performed to examine the differences between responses to stimuli for h1a/P1 (denoted by P1 in the Comparison column) and arc/P2 (denoted by P2 in the Comparison column). This analysis revealed that some differential responses to stimuli are only observed with a specific IEG/Presentation order channel.

| Region | Cell Type | Comparison | Estimate | Std. Error | z value | P adjusted |
| --- | --- | --- | --- | --- | --- | --- |
| **BLA** | Slc17a7 | Dom P1 - EC P1 | 0.434 | 0.208 | 2.081 | 0.059 |
| **BLA** | Slc17a7 | Dom P1 - Sub P1 | 0.198 | 0.03 | 6.713 | <0.001 |
| **BLA** | Slc17a7 | EC P1 - Sub P1 | -0.235 | 0.208 | -1.129 | 0.308 |
| **BLA** | Slc17a7 | Dom P2 - EC P2 | 0.572 | 0.209 | 2.733 | 0.011 |
| **BLA** | Slc17a7 | Dom P2 - Sub P2 | -0.126 | 0.03 | -4.266 | <0.001 |
| **EPd** | Slc17a7 | Dom P1 - EC P1 | 0.502 | 0.245 | 2.051 | 0.085 |
| **EPd** | Slc17a7 | Dom P1 - Sub P1 | 0.324 | 0.046 | 7.024 | <0.001 |
| **EPd** | Slc17a7 | EC P1 - Sub P1 | -0.178 | 0.245 | -0.727 | 0.526 |
| **EPd** | Slc17a7 | Dom P2 - EC P2 | 0.622 | 0.245 | 2.535 | 0.031 |
| **EPd** | Slc17a7 | Dom P2 - Sub P2 | 0.129 | 0.046 | 2.793 | 0.015 |
| **MeApd** | Slc32a | Dom P1 - EC P1 | 1.003 | 0.324 | 3.098 | 0.003 |
| **MeApd** | Slc32a | Dom P1 - Sub P1 | 0.485 | 0.035 | 13.842 | <0.001 |
| **MeApd** | Slc32a | EC P1 - Sub P1 | -0.518 | 0.324 | -1.599 | 0.136 |
| **MeApd** | Slc32a | Dom P2 - EC P2 | 0.501 | 0.323 | 1.552 | 0.147 |
| **MeApd** | Slc32a | Dom P2 - Sub P2 | 0.021 | 0.035 | 0.608 | 0.573 |
| **MeAav** | Slc32a | Dom P1 - EC P1 | 0.843 | 0.362 | 2.331 | 0.036 |
| **MeAav** | Slc32a | Dom P1 - Sub P1 | 0.414 | 0.049 | 8.513 | <0.001 |
| **MeAav** | Slc32a | EC P1 - Sub P1 | -0.43 | 0.362 | -1.188 | 0.295 |
| **MeAav** | Slc32a | Dom P2 - EC P2 | 0.66 | 0.361 | 1.829 | 0.107 |
| **MeAav** | Slc32a | Dom P2 - Sub P2 | 0.139 | 0.048 | 2.897 | 0.008 |
| **MeApv** | Slc17a7 | Dom P1 - EC P1 | 0.46 | 0.416 | 1.107 | 0.408 |
| **MeApv** | Slc17a7 | Dom P1 - Sub P1 | 0.085 | 0.127 | 0.668 | 0.604 |
| **MeApv** | Slc17a7 | EC P1 - Sub P1 | -0.375 | 0.407 | -0.922 | 0.483 |
| **MeApv** | Slc17a7 | Dom P2 - EC P2 | 0.145 | 0.412 | 0.353 | 0.79 |
| **MeApv** | Slc17a7 | Dom P2 - Sub P2 | -0.1 | 0.139 | -0.722 | 0.585 |
| **Ca3p** | Slc17a7 | Dom P1 - EC P1 | 0.387 | 0.222 | 1.746 | 0.129 |
| **Ca3p** | Slc17a7 | Dom P1 - Sub P1 | -0.136 | 0.031 | -4.363 | <0.001 |
| **Ca3p** | Slc17a7 | EC P1 - Sub P1 | -0.523 | 0.221 | -2.363 | 0.04 |
| **Ca3p** | Slc17a7 | Dom P2 - EC P2 | 0.238 | 0.221 | 1.073 | 0.332 |
| **Ca3p** | Slc17a7 | Dom P2 - Sub P2 | -0.17 | 0.031 | -5.475 | <0.001 |
| **Ca2Ca3d** | Slc17a7 | Dom P1 - EC P1 | 0.462 | 0.229 | 2.021 | 0.182 |
| **Ca2Ca3d** | Slc17a7 | Dom P1 - Sub P1 | 0.046 | 0.033 | 1.416 | 0.348 |
| **Ca2Ca3d** | Slc17a7 | Dom P1 - Dom P2 | 0.038 | 0.03 | 1.286 | 0.383 |
| **Ca2Ca3d** | Slc17a7 | EC P1 - Sub P1 | -0.416 | 0.229 | -1.821 | 0.247 |
| **Ca2Ca3d** | Slc17a7 | Dom P2 - EC P2 | 0.245 | 0.229 | 1.073 | 0.462 |
| **Ca2Ca3d** | Slc17a7 | Dom P2 - Sub P2 | -0.09 | 0.032 | -2.804 | 0.03 |

**Table S7.** Within-Subject IEG Responses in Each Stimulus Condition. Statistical results comparing the proportion of IEG positive cells of principal neuron-types in each brain region in response to P1 and P2.

| Region | Cell Type | Stimulus Condition | Comparison | Estimate | Std. Error | P Adjusted |
| --- | --- | --- | --- | --- | --- | --- |
| BLA | Slc17a7 | Dom-Sub | P2 - P1 | -0.4659 | -0.4659 | <0.001 |
| BLA | Slc17a7 | Sub-Dom | P2 - P1 | -0.3866 | -0.3866 | <0.001 |
| BLA | Slc17a7 | Dom-Dom | P2 - P1 | -0.4373 | -0.4373 | <0.001 |
| BLA | Slc17a7 | Sub-Sub | P2 - P1 | -0.1052 | -0.1052 | 5E-04 |
| BLA | Slc17a7 | EC-EC | P2 - P1 | -0.6461 | -0.6461 | <0.001 |
| EPd | Slc17a7 | Dom-Sub | P2 - P1 | -0.1309 | -0.2836 | 2E-04 |
| EPd | Slc17a7 | Sub-Dom | P2 - P1 | -0.1168 | 0.1755 | 1E-04 |
| EPd | Slc17a7 | Dom-Dom | P2 - P1 | 0.1461 | -0.0196 | <0.001 |
| EPd | Slc17a7 | Sub-Sub | P2 - P1 | 0.2458 | 0.1768 | <0.001 |
| EPd | Slc17a7 | EC-EC | P2 - P1 | 0.1788 | -0.1984 | <0.001 |
| MeAav | Slc32a | Dom-Sub | P2 - P1 | 0.2269 | -0.0315 | 0.0126 |
| MeAav | Slc32a | Sub-Dom | P2 - P1 | -0.3483 | 0.5334 | 0 |
| MeAav | Slc32a | Dom-Dom | P2 - P1 | 0.2237 | 0.1358 | 0.0097 |
| MeAav | Slc32a | Sub-Sub | P2 - P1 | 0.1607 | 0.412 | 0.0148 |
| MeAav | Slc32a | EC-EC | P2 - P1 | 0.2182 | 0.31 | 0.0113 |
| MeApd | Slc32a | Dom-Sub | P2 - P1 | 0.0598 | 0.0808 | 0.066 |
| MeApd | Slc32a | Sub-Dom | P2 - P1 | -0.2242 | 0.5881 | <0.001 |
| MeApd | Slc32a | Dom-Dom | P2 - P1 | 0.1448 | 0.0833 | <0.001 |
| MeApd | Slc32a | Sub-Sub | P2 - P1 | 0.1383 | 0.5429 | <0.001 |
| MeApd | Slc32a | EC-EC | P2 - P1 | 0.1585 | 0.5926 | <0.001 |
| MeApv | Slc17a7 | Sub-Dom | P2 - P1 | -0.2836 | -0.6802 | <0.001 |
| MeApv | Slc17a7 | EC-EC | P2 - P1 | 0.1755 | -0.457 | 4E-04 |
| Ca2Ca3d | Slc17a7 | Dom-Sub | P2 - P1 | -0.0196 | -0.1309 | 0.6426 |
| Ca2Ca3d | Slc17a7 | Sub-Dom | P2 - P1 | 0.1768 | -0.1168 | 1E-04 |
| Ca2Ca3d | Slc17a7 | Dom-Dom | P2 - P1 | -0.1984 | 0.1461 | 0.0017 |
| Ca2Ca3d | Slc17a7 | Sub-Sub | P2 - P1 | -0.0315 | 0.2458 | 0.5561 |
| Ca2Ca3d | Slc17a7 | EC-EC | P2 - P1 | 0.5334 | 0.1788 | <0.001 |
| Ca3p | Slc17a7 | Dom-Sub | P2 - P1 | 0.1358 | 0.0598 | 0.0076 |
| Ca3p | Slc17a7 | Sub-Dom | P2 - P1 | 0.412 | -0.2242 | <0.001 |
| Ca3p | Slc17a7 | Dom-Dom | P2 - P1 | 0.31 | 0.1448 | <0.001 |
| Ca3p | Slc17a7 | Sub-Sub | P2 - P1 | 0.0808 | 0.1383 | 0.0359 |
| Ca3p | Slc17a7 | EC-EC | P2 - P1 | 0.5881 | 0.1585 | <0.001 |

